# The impact of phosphorylated Pten at threonine 366 on cortical connectivity and behaviour

**DOI:** 10.1101/2021.09.22.461341

**Authors:** Julia Ledderose, Jorge A. Benitez, Amanda J. Roberts, Rachel Reed, Willem Bintig, Robert Sachdev, Frank Furnari, Britta J. Eickholt

## Abstract

The lipid phosphatase Pten (phosphatase and tensin homologue on chromosome 10) is a key tumour suppressor gene and an important regulator of neuronal signalling. Pten mutations have been identified in patients with autism spectrum disorders, characterized by macrocephaly, impaired social interactions and communication, repetitive behaviour, intellectual disability, and epilepsy. Pten enzymatic activity is regulated by a cluster of phosphorylation sites at the C-terminus of the protein. Here we specifically focussed on the role of Pten T366 phosphorylation and generated a knock-in mouse line in which Pten T366 was substituted with alanine (*Pten^T366A/T366A^*). We identify that phosphorylation of Pten at T366 controls neuron size and connectivity of brain circuits involved in sensory processing. We show in behavioural tests that *Pten^T366/T366A^* mice exhibit cognitive deficits and selective sensory impairments, with significant differences in male individuals. We identify restricted cellular overgrowth of cortical neurons in *Pten^T366A/T366A^* brains, linked to increases in both dendritic arborization and soma size. In a combinatorial approach of anterograde and retrograde monosynaptic tracing using rabies virus, we characterize differences in connectivity to the primary somatosensory cortex of *Pten^T366A/T366A^* brains, with imbalances in long-range cortico-cortical input to neurons. We conclude that phosphorylation of Pten at T366 controls neuron size and connectivity of brain circuits involved in sensory processing and propose that PTEN T366 signalling may account for a subset of autism-related functions of Pten.

## Introduction

Mutations in the PTEN gene have been linked to patients with autism spectrum disorders (ASD) ^1–6^. ASD is a neurodevelopmental disorder with mild to severe symptoms in motor skills or in responses to sensory stimuli in young patients, which result in social and communication challenges in adults ^7,8^. PTEN-ASD patients show neuroanatomical abnormalities, increase in white matter and cortex, and enlargements of perivascular spaces ^9–11^. Cognitive deficits in these patients may be related to changes in sensory processing, neuronal connections or cortical circuits ^12^.

Autism spectrum disorders is a highly inheritable disease with more than 35 ASD missense mutations identified in the PTEN gene, often associated with macrocephaly ^4,5^. The macrocephaly phenotype observed in ASD patients carrying PTEN mutations correlates with findings from loss-of-function mouse models ^2,13–15^. In this context, -- since homozygous Pten mice are lethal -- Pten conditional knock-out (KO) mice have been analyzed to understand the function of Pten during brain development. These concerted efforts shed light on several ASD-linked phenotypes ^2,13,16^. Germline haploinsufficient Pten mice (*Pten^+/−^*), an extensively studied mouse ASD model, exhibit impaired social behaviour in connection to neuronal hypertrophy and increased brain size ^16–18^. Also, brain volume in *Pten^+/^* mice scales differently during development compared to wildtype (wt) mice ^13^, revealing another Pten phenotype typical of ASD.

Pten is a regulator of the PI3K signalling pathway^19^, and is involved in neuronal growth ^20–22^. Loss of Pten in mice leads to neuronal hypertrophy and to changes in morphology ^23,24^. Pten-deficient neurons show increased dendritic arborisation and changes in their connectivity ^15,25–27^. In *Pten^fl/fl^ Nse-Cre* KO mice, there is an increase in dendritic arborization and spine density of the hippocampal CA3 neurons, and a collective increase in the axons in the dentate gyrus ^2^. When Pten is deleted in cortical neurons (*Camk2a-cre^+/−^*; *Pten^loxP/loxP^*; *Thy1-gfp*), pyramidal layer (L) 2/3 neurons show a specific increase in dendritic arborization and spine density ^25^. Pten works primarily at the plasma membrane, dependent on cytosolic AKT/mTOR signaling ^23,24^, however it also has PI3K-independent, and lipid phosphatase-independent signalling mechanisms that occur at the level of the nucleus ^28–30^. Furthermore, studies have suggested that dysregulation of mTOR signalling could contribute to the development of ASD ^31,32^.

The regulation of PTEN activity and function primarily involves phosphorylation of a cluster of serine and threonine residues in the Pten C-terminus. Several phosphorylation sites at the C-terminal tail of the Pten gene are required for regulation of Pten stability and activity ^19^. The phosphorylation site at T366 can be phosphorylated by polo-like-kinase (Plk3) and glycogen synthase kinase 3 (GSK3), leading to either stabilization of the PTEN protein in glioma cells ^33,34^ or to destabilization in epithelial cells ^35–37^.

We generated mice, in which the normal phosphorylation at T366 was abrogated through substitution with alanine (*Pten^T366A/T366A^*). These mice develop sensory abnormalities in exploration, freezing, and in their response to vibrissal stimuli, and their brains show changes in cortical connectivity. We conclude that intact phosphorylation at Pten T366 is essential for the establishment of proper cortical connectivity and for normal sensory processing. Our work identifies that T366 phosphorylation of PTEN might serve as important signalling regulator in ASD.

## Methods

### Ethics statement and handling of mice

All experiments were conducted under the licenses T0143/11, G0621/12 and G0189/14 in accordance with the guidelines of the Universitätsmedizin Charité, Berlin, and the local authorities, the Landesamt für Gesundheit und Soziales (LaGeSo). Behavioural studies were conducted in the Animal Models Core at The Scripps Research Institute and were approved by The Scripps Research Institute’s Institutional Animal Care and Use Committee, and met the NIH guidelines detailed in the ‘Guide for the Care and Use of Laboratory Animals’. Transgenic knock-in *Pten^T366A/T366A^* mice in an FVB/N background and FVB/N wt mice for control experiments were used. For the behavioural experiments, *Pten^T366A/T366A^* mice crossed with wt mice of C57BL6/N background and C57BL6/N mice for control experiments were used. All mice were kept in groups of two to three individuals under standard conditions in a 12-hour day-night-cycle with water and food available ad libitum.

### Generation of *Pten^T366A/T366A^* knock-in mice

A BAC construct containing the mouse *Pten* gene was used as a genomic source to create a targeting vector. Briefly, a 5’ homology arm of exon 8 followed by exon 9 containing a threonine 366 to alanine mutation and a right homology arm containing intron 9 were inserted into targeting vector pKO V915 (Dana Farber Cancer Center) flanking a LoxP-pGK-Neo-LoxP selection cassette. The linearized targeting vector was electroporated into 129S6/SV EV Tac embryonic stem cells and clones were screened by southern blot analysis for correct targeting into the *Pten* locus. Two correctly targeted ES cell clones were injected into blastocysts to generate chimeric mice in C57BL/6J background. Mice with greater than 80 % chimerism, as determined by agouti coat color, were bred with C57BL/6J mice and progeny were screened by southern blot to ascertain germline transmission. Heterozygous mice were bred with E2a-cre mice (Jackson Lab, #003314) to excise the pGK-Neo selection cassette and progeny were screened by PCR to confirm successful cre-mediated excision. Heterozygous mice lacking pGK-Neo were bred with C57BL/6J mice to generate *Pten^T366A/T366A^* mice.

### DNA isolation, genotyping, and sequencing

DNA from tail or ear tissue of *Pten^T366A/T366A^* mice was incubated in 80% Proteinase-K in TE buffer for 6 hours. DNA was purified with a DNA purification kit (Thermo Fisher Scientific) and analyzed with standard PCR conditions. The genotype of *Pten^T366A/T366A^* mice was determined by PCR with sequencing of the PCR product. Primers for targeting the Pten gene location at threonine 366 were forward (5’-AGCAGTGCCCTTCAGAATTC-3’), and reverse (5’-TCAGCCACTTCAGCTGGTGAC-3’), resulting in a PCR product of 600 bp. Sequencing was performed with the forward primer.

### Primary cell culture

Mouse hippocampi and cortices were dissected from wt FVB/N and *Pten^T366A/T366A^* mice at embryonic stage (E) 16.5 and dissociated in papain (Worthington) according to the manufacturer’s protocol. 24-well dishes (MatTek Corporation) were coated with 30 ng/μl poly-D-lysine and hippocampal cells and cortical cells were plated in growth medium (Neurobasal medium supplemented with 1 % Glutamax, 1 % B27). Cells were fixed in 4 % paraformaldehyde (PFA) after 2, 4, 8 and 14 days *in vitro* (DIV) and used for immunohistochemistry.

### Protein lysate preparation and western blotting

Whole mouse brains at postnatal day (P) 10 were weighed and homogenized in 4 × the volume of RIPA buffer (50 mM Tris-HCl, pH 7.4, 150 mM NaCl, 0.5% sodium deoxycholate, 1 % NP40, 0.1% sodium dodecyl sulphate) supplemented with protease inhibitors (Calbiochem set III) and phosphatase inhibitors (1 mM Na_2_MO_4_, 1 mM NaF, 20 mM β-glycerophosphate, 1 mM Na_3_VO_4_, 500 nM cantharidin) using a glass pistil. Homogenates were centrifuged at 20,000 × g and the supernatant was collected for further protein quantification analysis using BCA Thermo Scientific Pierce™ Protein Assay. The neurons were washed once with cold PBS and lysed in cold RIPA buffer (including protease and phosphatase inhibitors). Cell lysates were centrifuged at 20,000 × g and the supernatant was transferred to a tube containing Roti load I sample buffer. Western blot analysis was performed as previously described ^38^. Protein lysates in Roti-Load sample buffer (5 to 10 μg total protein) were loaded on 8 % SDS gels, stacked at 60 V for 30 min, and separated at 120 V for 60 min. Proteins were transferred to a nitrocellulose membrane for 2 hours in a wet blot tank system (Bio-Rad), and membranes were blocked with 5% skim milk for 30 min at room temperature before incubating with primary antibody. Primary antibodies were prepared (1:1000) in 5 % skim milk and incubated overnight at 4°C. Following incubation, membranes were washed three times with TBS-T at room temperature for 5 min. Secondary antibodies were prepared (1:3000) in 5 % skim milk, and membranes incubated for 30 min at room temperature, washed three times with TBS-T and imaged using the Fusion SL system from Vilber Lourmat. Quantification of band densities was performed by Fiji/ ImageJ. The area of the band and the mean grey value were measured to obtain a relative density. For relative quantifications, measurements were normalized to loading control.

### Antibodies

#### Western blotting

Primary antibodies: anti-PTEN rabbit (Cell Signalling, #9559), anti-pThr308AKT rabbit (Cell Signalling, #9275), anti-pSer473AKT rabbit (Cell Signalling, #4060), anti-pGSK3b mouse (BD Transduction Laboratory, #610202), anti-Tubulin-beta III mouse (Covance #PRB-435P). Secondary antibodies HRP conjugated anti-rabbit, and anti-mouse (Vector Labs, #PI1000, catalog no. PI2000)

#### Immunohistochemistry

Primary antibodies: anti-MAP2 mouse (1:500, Sigma M9942), anti-MAP2 guinea pig (1:500, Synaptic System, 188 004), anti-GFP chicken (1:5000, Abcam ab13970), anti-BrdU rat (1:250, Biorad, OBT0030), anti-Cre rabbit (1:1000, Abcam ab41104), anti-FoxP2 (1:1000, Abcam ab16046), anti-NeuN mouse (1:1000, Millipore MAB377), anti-Cux1 rabbit (1:1000, Santa Cruz, sc13024), anti-CTIP2 rat (1:1000, Abcam ab18465). Secondary antibodies (all 1:500), anti-mouse Alexa Flour 488 (Dianova, 015-540-003), anti-mouse Cy5 (Dianova, 715-175-151), anti-chicken Alexa Flour 488 (Dianova, 703-546-155).

### Production of recombinant adenovirus and rabies virus

Viruses were produced in the Viral Core facility of the Universitätsmedizin Charité, Berlin (www.vgf.charite.de) ^39–41^using standard protocols ^42^. An adenovirus, tagged to a nuclear green fluorescent protein (AAV-nls-GFP) was used for morphological analysis. For combined anterograde tracing with monosynaptic rabies virus tracing, three different viruses were used. We first injected an AAV1.hSyn.Cre.WPRE.hGH (Addgene, #105553) targeting thalamic POm and VPm region. After three weeks of Cre expression, we injected an AAV virus coupled to a nuclear GFP and an EnvA interacting TVA receptor, and to a rabies G protein (AAV-EnvA-TvA-GFP) to initiate virus expression for the rabies virus. The subsequently injected rabies virus was coupled to a mCherry fluorophore and contained a deleted rabies glycoprotein (RABB-SADB19dG-mCherry)^41^.

### Injections in new-born mice

Viral injections into new-born mice were carried out as described ^43^. Newborn pups postnatal day (P) 0 were removed from their mother and briefly anaesthetized by isoflurane. AAV-GAG-nls-GFP virus was injected into both lateral ventricles using a 34-gauge needle and a microprocessor infusion pump with an injection speed of 125 nl/sec. After injection, pups were allowed to recover on a warming plate at 37 °C, and replaced in the home cage with their mother.

### Stereotaxic injections in adult mice

Before surgery, mice were anaesthetized with ketamine (100 mg/kg) and xylazine (10 mg/kg) and head-fixed on a stereotaxic frame with non-puncture ear bars and a nose clamp (Kopf Stereotaxic device, California, USA, Inc.). The extend of the anesthetic was confirmed by a toe pinch. For analgesics, Carpofen (5 mg/kg) was injected intraperitoneal, and lidocaine was injected under the skull before surgery. The scull was opened with scissors and a craniotomy (~1 mm) was made on the left hemisphere above the injections sites. The following coordinates were used for somatosensory cortex (S1, AP −1.5, lateral: 2.3 to 2.7), and for POm/ VPm thalamus (AP, 1.5, lateral, 1.5, ventral 3.0). Negative pressure was applied to tip-fill the fine glass pipette with a virus solution. We first injected 200 nl of virus under constant positive pressure with a flow rate of 20 nl/min (QSI, Quintessential Stereotaxic Injector). During surgery, the brain tissue was kept moist by applying sterile PBS at regular intervals, the eyes were lubricated with Bepanthen eye cream (Bayer). Following stereotaxic injection, the craniotomy was cleaned, the skull sutured, and the mouse returned to its home cage for recovery. Caprofen (5 mg/kg) was administered for two days after surgery to reduce postoperative pain.

### BrDU proliferation experiments

Birth dating experiments with BrdU were carried out as described previously ^44^. Time-pregnant female mice were injected intraperitoneally with 50 mg/kg BrdU (5-bromo-2’-deoxyuridine, BrdU; Accurate Chemical & Scientific Corporation) at E11.5, E13.5 and E15.5. BrdU was expressed until P1 and P8. For analysis, the number of BrdU positive neurons were analyzed in somatosensory cortices of new-born P1 mice and in somatosensory cortices of P8 mice.

### Tissue processing and immunohistochemistry

Neuronal cultures were fixed in 4 % PFA for 30 min. Neurons were permeabilized in PHEM with 0.1 % Triton for 10 min and blocked in PHEM with 4 % goat serum. Primary antibodies were incubated for 1 hour followed by PBS washes and incubation of secondary antibodies for 1 hour.

Mice were deeply anaesthetized with isoflurane and trans-cardially perfused with PBS and 4 % PFA, followed by fixation in 4 % PFA overnight. Brains were soaked in 30 % sucrose overnight and cut with a microtome (70 μm to 100 μm). Brain sections were stored in PBS for immediate immunohistochemistry and in cryo-protection medium (Ethylene ethanol 30 %, Glycerol 30 % in PBS) for long-term storage. For immunohistochemistry, brain sections were placed in blocking solution (5 % NGS, 0.1 % Triton X-100 in PBS) for 2 hours and incubated in primary antibodies overnight at room temperature. Secondary antibodies were applied following washes in PBS. Brain sections were counterstained with HOECHST (1:5000, Sigma), washed with PBS and mounted in glycerol (80 %, 2.5 % DAPCO in PBS).

### Nissl staining

Brain sections were cut with a microtome and mounted on Super Frost Plus Slides (Thermo Fisher Scientific). Sections were incubated in Nissl solution for 40 s to 1 min, then washed three times in water, and dehydrated in ethanol (50 % EtOH, 70 % EtOH, 80 % EtOH, 95 % EtOH, 5 min each step), followed by incubation in Xylol. Slices were mounted in Eukitt-quick-hardening mounting medium (Fluka) and stored at 4 °C.

### Behavioural experiments

Two cohorts of 12 to 16 week old mice were used in the behavioural experiments (cohort 1: 8 male wt, 5 male *Pten^T366A/T366A^*, 6 female wt, 5 female *Pten^T366A/T366A^;* cohort 2: 5 male wt, 5 male *Pten^T366A/T366A^*, 6 female wt, 5 female *Pten^T366A/T366A^*). Cohort 1 was tested in the Y maze, open field, Barnes maze and conditioned fear tests. Cohort 2 was tested in the Y maze, hanging wire, rotarod, von Frey, Morris water maze, conditioned fear and vibrissae-stimulated reflex tests. All tests were performed during the dark (active) phase of the circadian rhythm with 4 to 7 intervening days. Data from the cohorts were combined in the case of replicated tests.

#### Y maze test

Spontaneous alternation behaviour, a measure of spatial working memory, exploratory behaviour, and responsiveness to novelty ^45,46^, was tested using a Y maze with 34 × 8 × 14 cm arms. Each mouse was tested in a single 5-min trial and spontaneous alternations, sets of three unique arm choices, were recorded. Because mice have the opportunity to do perform repeated entries into a single arm, there is a chance performance level of 22 % (2/9) for spontaneous alternations ^47,48^.

#### Hanging wire test

The hanging wire test allows for the assessment of grip strength and motor coordination ^49,50^. Mice were held so that only their forelimbs contact an elevated metal bar (2 mm diameter, 45 cm long, 37 cm above the floor) held parallel to the table by a large ring stand and let go to hang. Each mouse was given three trials separated by 30 s. Each trial was scored as follows and the average for each mouse was calculated: 0 — fell off, 1 — hung onto the wire by two forepaws, 2 — hung onto the wire by two forepaws, but also attempted to climb onto the wire, 3 — hung onto the wire by two forepaws plus one or both hindpaws around the wire, 4 — hung onto the wire by all four paws plus tail wrapped, 5 — escaped (crawled to the ring stand and righted itself or climbed down the stand to the table). Latency to falling off was measured up to a maximum of 30 s.

#### Rotarod test

Rotarod balancing requires a variety of proprioceptive, vestibular, and finetuned motor abilities as well as motor learning capabilities ^51^. A Roto-rod Series 8 apparatus (IITC Life Sciences, Woodland Hills, CA) was used which records test results when the animal drops onto the individual sensing platforms below the rotating rod. An accelerating test strategy was used whereby the rod started at 0 rpm and then accelerated to 10 rpm. The mice were tested in two sets of three trials per day for four days, for a total of 24 trials.

#### Open field test

This test predicts how animals respond when introduced into a brightly illuminated open arena ^52^. It is a classic test of “emotionality” used to measure anxiety-like responses of rodents exposed to stressful environmental stimuli (brightly illuminated open spaces) and to capture spontaneous activity measures. The apparatus is a square white Plexiglas (50 x 50 cm) open field illuminated to 600 lux in the center. Each animal is placed in the center of the field and several behavioural parameters (distance traveled, velocity, center time, frequency in center) are recorded during a 10-min observation period and analyzed using Noldus Ethovision XT software. Time spent grooming was also assessed.

#### von Frey test

Mechanical sensitivity was assessed by the application of von Frey filaments of varying forces (0.16, 0.4, 1, 2, 4, 6, 8 g) perpendicularly to the hind paw ^53^. If the mouse withdrew its paw, a positive response was recorded. Each filament was tested a total of 10 times.

#### Vibrissae-stimulated reflex test

This is a test of sensorimotor function. Mice are held by their torso while their vibrissae are brushed along a tabletop. In normal mice, this elicits the placement of a forelimb, ipsilateral to the stimulation side, on the table ^54^. Ten trials per side were performed on two days, the number of successful placements for each mouse was tabulated and averaged.

#### Barnes maze test

This is a spatial memory test ^55–57^ sensitive to impaired hippocampal function ^58^. Mice learn to find an escape chamber (19 x 8 x 7 cm) below 1 of 20 holes (5 cm diameter, 5 cm from perimeter) below an elevated brightly lit and noisy platform (75 cm diameter, elevated 58 cm above floor) using cues placed around the room. Spatial learning and memory were assessed across 4 trials (maximum time was 3 min) and then directly analyzed on the final (fifth) probe test in which the tunnel was removed and the time spent in each quadrant was determined and the percent time spent in the target quadrant (the one originally containing the escape box) was compared with the average percent time in the other three quadrants. This is a direct test of spatial memory as there is no potential for local cues to be used in the mouse’s behavioural decision.

#### Morris water maze test

The water maze test is used to assess spatial learning and memory in rodents (Morris, 1981; Puzzo et al., 2014). Mice are placed into a circular tub filled with opaque water and they learn over repeated trials to locate a hidden platform onto which they can sit and escape from the swimming. Each animal underwent two trials per day for four days, with a fixed platform location, and a random start position. After being released into the water, each animal was allowed to swim until the platform is found or 90 s had elapsed, at which point the experimenter gently guided the mouse to the platform. A probe trial was given after the completion of training (day 5), in which the platform was removed from the water maze and the animal was allowed to swim freely for 90 s. The amount of time spent in each quadrant, the number of times a mouse entered a quadrant, and the number of times the mouse crossed the platform location (annulus crossings) were recorded using Noldus Ethovision software.

#### Conditioned fear test

In this procedure, mice learn to associate a novel environment (context) and a previously neutral stimulus (conditioned stimulus, a tone) with an aversive foot shock stimulus ^60^. It allows the assessment of both hippocampus-dependent and amygdala-dependent learning processes in the same mouse ^61,62^. Conditioned animals, when exposed to the conditioned stimuli, tend to refrain from all but respiratory movements by freezing. Freezing responses can be triggered by exposure to either the context in which the shock was received (context test) or the conditioned stimulus (CS+ test). Briefly, mice were habituated to the system (Freeze Monitor, Med Associates, VT) to measure baseline freezing behaviour on day 1 (5 min trial) and then on day 2 were conditioned with two 0.6 mA foot shocks given in the final 2 s of cue exposure (30 s, 3000 Hz, 80 dB sound + white light) in a 6 min trial. On day 3, contextual conditioning (as determined by freezing behaviour) was measured in a 5 min trial in the chamber where the mice were trained (context test). The following day, the mice were tested for cued conditioning (CS+ test). The mice were placed in a novel context for 3 min, after which they were exposed to the conditioned stimuli (light + tone) for 3 min. For this test, the chamber was disguised with new walls (white opaque plastic creating a circular compartment in contrast to a clear plastic square compartment) and a new floor (white opaque plastic in contrast to metal grid). Freezing behaviour (i.e., the absence of all voluntary movements except breathing) was measured in all sessions by real-time digital video recordings calibrated to distinguish between subtle movements, such as whisker twitches, tail flicks, and freezing behaviour. Freezing behaviour is indicative of the formation of an association between the particular stimulus (either the environment or the tone) and the shock; i.e. that learning has occurred.

### Confocal imaging

Images were taken on confocal laser scanning microscopes (Leica SP5, Leica SP8 Nikon A1Rsi+) with 10x air, and a 20x or a 63x oil immersion objective (Leica: HCX PL APO 20x/0.7, HCX PL APO 63x/1.20 W motCORR CS; Nikon 20 x Plan Apo, Air 0.8 NA, 1.000 DIC N2 VC). A 405 nm laser was used for detection of fast blue, a 561 nm laser for detection of tdTom or mCherry, and a 488 nm laser for detection of GFP (Leica, 425/70, 515/25, 590/70; Nikon, 450/50, 525/50, 595/50).

### Data analysis

#### Analysis of soma size and dendritic arborization of neurons in S1 cortex

For the analysis of somata, maximal intensity projections of confocal images were used, and soma size was measured with ImageJ/Fiji. The maximal projection was converted to an 8-bit black and white image, and the threshold was adjusted manually. For analysis of dendritic arborisation, maximal intensity projections of confocal images were used and dendrites reconstructed with the imaging analysis software IMARIS. The complexity of the dendrites was analyzed with Sholl analysis ^63^. L2/3 pyramidal neurons were defined by a triangular soma shape and by having one apical dendrite reaching L1, as well as basal and apical dendrites. L4 interneurons, spiny stellate neurons, were defined by having a local arborisation, and somata having a round shape.

#### Defining cortical layers

We defined cortical areas and cortical layers according to the Allen brain reference atlas (https://mouse.brain-map.org/) and to the Paxinos Atlas. For cortical layers, we used the following definitions: S1, from pia: L1, 100 μm, L2/3, 100 μm to 300 μm; L4: 300 μm to 400 μm; L5a, 400 μm to 500 μm; L5b, 500 μm to 700 μm; L6a, 700 μm to 850 μm; L6b, 850 μm to 950 μm or 100 μm from white matter.

#### Statistical analysis

All data were statistically analyzed with Graphpad/ Prism software. Statistics for behaviour, soma size and dendritic lengths and connectivity were calculated by Students-t-test or two-way ANOVA, followed by Bonferroni posthoc test. The Sholl analysis was analyzed by two-way-ANOVA. All data are shown as mean ± S.E.M. (standard error), p-values are indicated as: *p<0.05, **p<0.01, ***p<0.001.

#### Data availability

Data are available from the corresponding author upon request.

## Results

### Generation of Pten T366 deficient (*Pten^T366A/T366A^*) mice

To test the contribution of Pten T366 to brain development and function, we analyzed mice in which Pten T366 phosphorylation was inactivated by the introduction of a threonine (T) to alanine (A) amino acid substitution, leading to constitutive inactive phosphorylation at PTEN T366 (thereafter abbreviated as T366A). Homozygous *Pten^T366A/T366A^* mice are viable and fertile. Embryos were born at expected Mendelian ratios and did not display overt phenotypes. In experiments for cortical development, we used heterozygous *Pten^T366A/+^* crosses to compare *Pten^T366A/T366A^* with wt siblings from the same litter. Forebrain homogenates obtained from *Pten^T366A/T366A^* mice showed normal expression levels of PTEN protein when compared to wt samples. Similarly, prominent signalling effectors of Pten activity such as pAkt, pS6, and pGSK3β did not show any differences in lysates (**Supplementary Fig. 1**). These results show that systemic loss of Pten T366 phosphorylation does not overtly affect normal embryonal development, PTEN protein stability or PTEN signalling pathways.

### *Pten^T366A/T366A^* mice show selective sensory impairments

Studies have shown a reduction in social learning and social interaction in Pten neuron-specific enolase (Nse) promoter-driven cre transgenic mice^2^. Heterozygous *Pten^+/−^* mice show increased anxiety and hyperactivity under stressful environments and decreased attention^16,17^, and a depression-like-phenotype in *Pten^+/−^* male mice ^18^. To characterize whether the loss of Pten phosphorylation at T366 leads to changes in sensory motor behaviours, we performed standard behavioural tests in *Pten^T366A/T366A^* mice (**Figure 1, Supplementary Table 1**). Exploratory behaviour in the open field was similar in *Pten^T366A/T366A^* mice compared to their wt littermates, in terms of both the distance travelled and time spent in the center of the field (**Figure 1A**). Motor performance and strength as determined using the hanging wire test (**Figure 1B**) and the rotarod (**Figure 1C**) were similar in *Pten^T366A/T366A^* mice and their wt siblings. Response to mechanical stimulation in the Von Frey test also did not differ between genotypes (data not shown).

**Figure 1.**
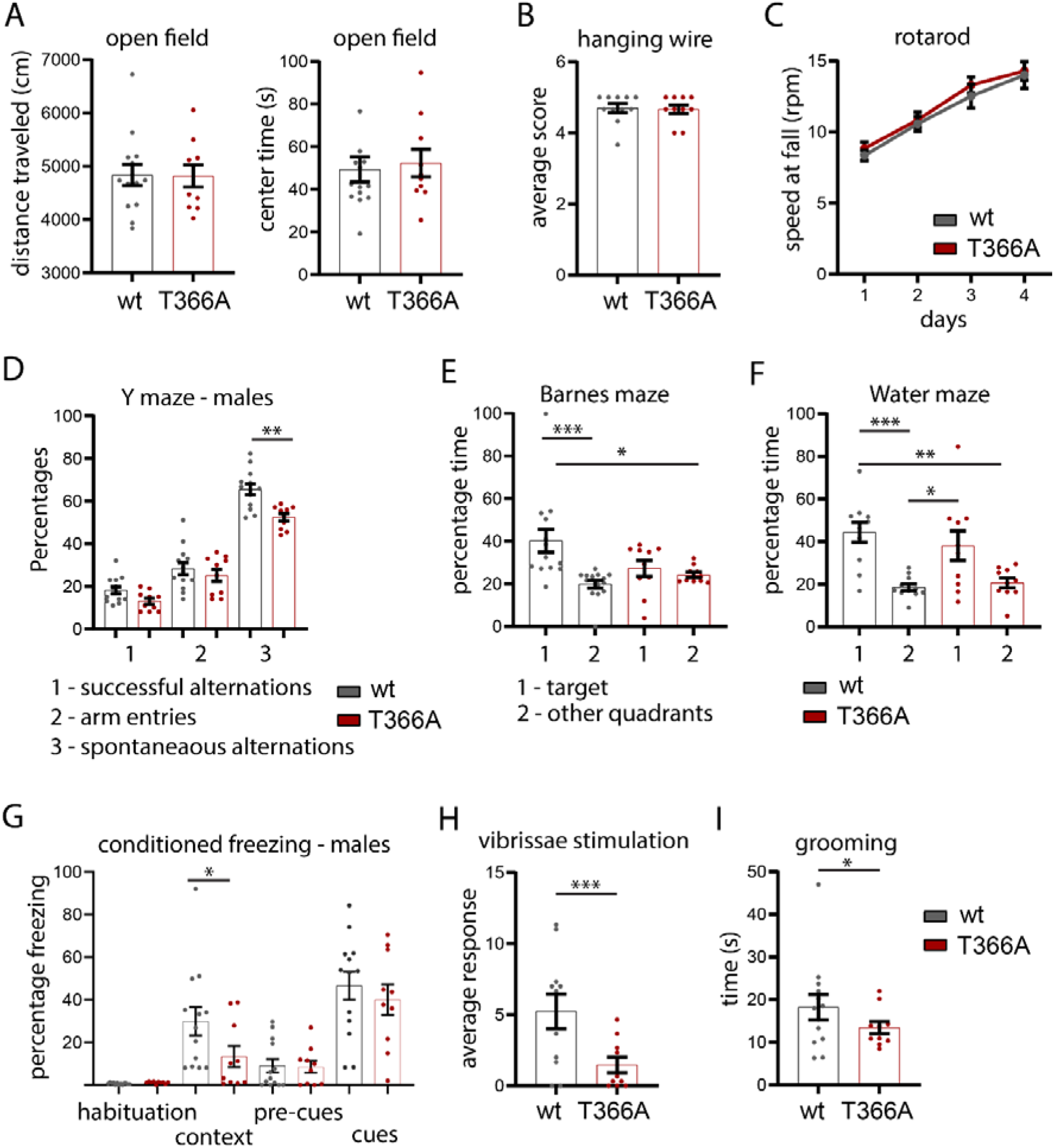
Behaviour in *Pten^T366A/T366A^* mice. (**A**) *Pten^T366A/T366A^* mice and wt mice tested for general exploratory behaviour in the open field. (**B, C**) Motor skills as determined in the in hanging wire test (**B**) and in the rotarod (**C**). (**D**) Percentage of spontaneous alterations in the Y maze in male *Pten^T366A/T366A^* and wt mice. (**E**) Percentage of time that *Pten^T366A/T366A^* and wt mice spent in the target area and other quadrants of the Barnes maze. (**F)** Percentage of time that *Pten^T366A/T366A^* and wt mice spent in the target area and other quadrants of the Morris water maze. (**G)** Percentage freezing in the conditioned fear-induced freezing in male *Pten^T366A/T366A^* and wt mice. (**H**) The average number of responses in the vibrissae-stimulated reflex test of *Pten^T366A/T366A^* and wt sibling. (**I**) Grooming time in *Pten^T366A/T366A^* and wt siblings. Statistical analysis with two-way ANOVA (Bonferroni post hoc test), unpaired t-test, Wilcoxon-test (grooming), *p<0.05; **p<0.01, ***p<0.001. For analysis details see **Supplementary Table 1.**

There were, however, differences in behavioural testing for cognitive abilities in *Pten^T366A/T366A^* mice. In the Y maze, male *Pten^T366A/T366A^* mice showed a decrease in spontaneous alternation behaviour relative to wt siblings (**Figure 1D**). The performance of *Pten^T366A/T366A^* mice, when both sexes were evaluated in the Barnes maze (**Figure 1E**), and the Water maze was diminished (**Figure 1F**). *Pten^T366A/T366A^* mice also performed poorly (compared to wt mice) in the probe tests, suggesting that these mice have spatial memory deficits. In the conditioned fear test, male *Pten^T366A/T366A^* mice showed less freezing in the context task compared to their wt siblings (**Figure 1G**), supporting that decreased hippocampus-mediated contextual learning and memory occurs in these mice.

*Pten^T366A/T366A^* mice also demonstrated a decreased number of responses to vibrissae stimulation (**Figure 1H**) and grooming time in the open field test was reduced (**Figure 1I**). Overall, these phenotypes tend towards differences in cognitive and sensory processing in *Pten^T366A/T366A^* mice, which could be based on differences of cortical morphology or connectivity of cortical neurons.

### Layer formation and cortical proliferation are normal in *Pten^T366A/T366A^* brains

Germline Pten mutations show disrupted cortical proliferation ^64^, and affected scaling across brain areas during development ^13^. To characterize whether loss of Pten T366 phosphorylation alters cortical layering, we performed immunolabeling, exploiting different layer specific markers. We used immunolabeling against NeuN for all neurons, and Cux1 for cortical L2/3 and L4, Ctip2 for cortical L5, and FoxP2 for cortical L6 (**Figure 2A-2D, Supplementary Table 2**). The overall percentage of NeuN neurons in the single layers did not show significant differences in *Pten^T366A/T366A^* mice compared to wt mice at P14 (**Figure 2A**). Neither the percentage of Cux1 positive neurons in L2/3 (**Figure 2B**), nor the percentage of Ctip2 positive neurons in L5 (**Figure 2C**) or the percentage of FoxP2 positive neurons in L6 (**Figure 2D**) differed significantly between the genotypes when analysed with relation to HOECHST counterstain. This analysis suggests that cortical layers develop normally in *Pten^T366A/T366A^* mice.

**Figure 2.**
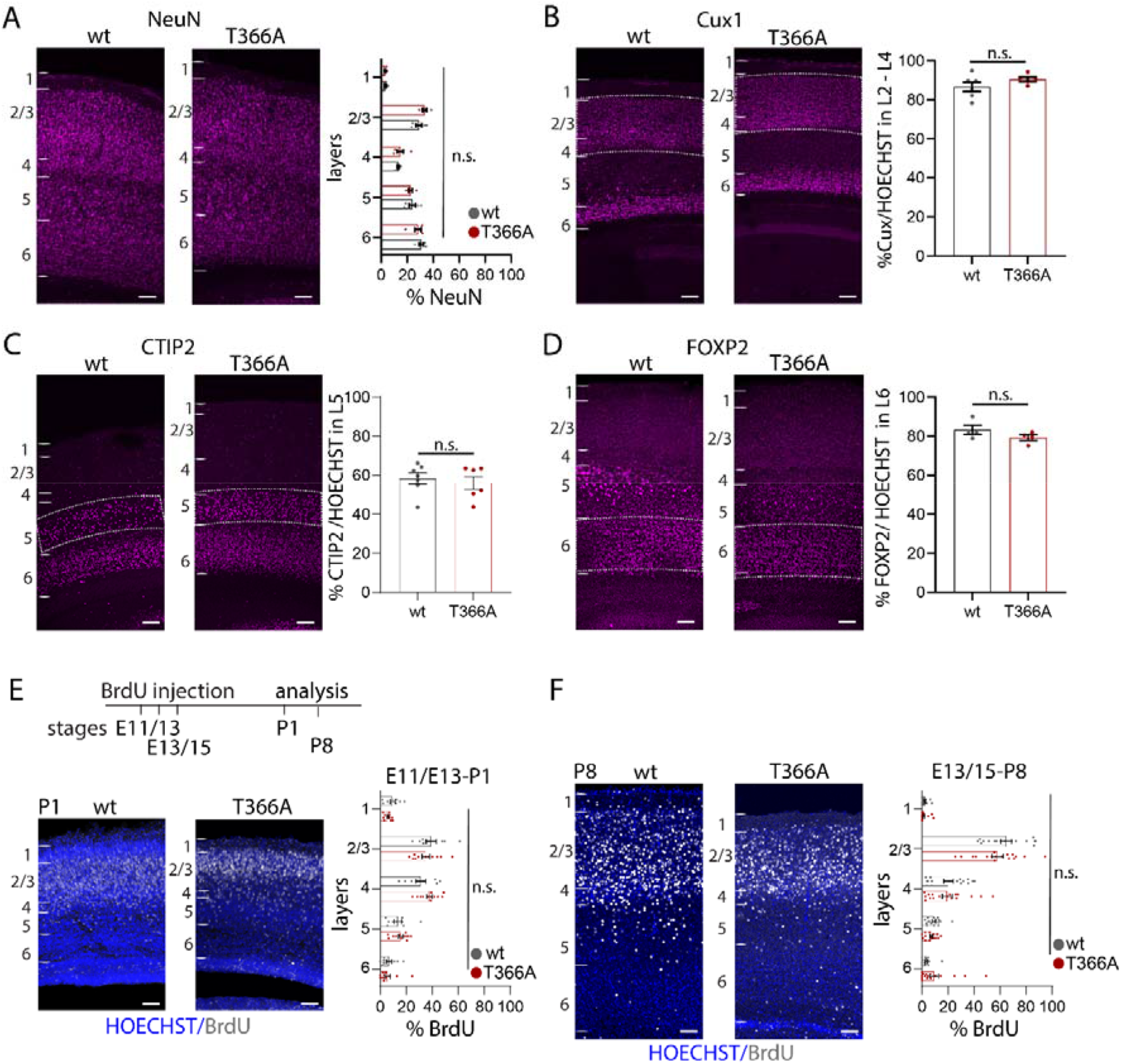
Cortical layers and proliferation in *Pten^T366A/T366A^* brains. (**A-D**) Immunolabeling for cortical layer markers and the percentages of neurons in *Pten^T366A/T366A^* and wt mice. (**A-D**) The percentage of (**A**) NeuN positive neurons in L1 to L6. (**B-E**) The percentage of **(B)** Cux1 positive neurons in L2 to L4 (**C**) Ctip2 positive neurons in L5 (**D**) FoxP2 positive neurons in L6 in relation to all neurons counterstained with HOECHST. **(E)** Proliferation of cortical progenitor cells in *Pten^T366A/T366A^* and wt mice. Schematic for experimental set-up, BrdU injections were performed in time-pregnant mice at E11.5, E13.5 and E15.5. Analysis was undertaken at P1 and P8. Example images of BrdU labelling and HOECHST counterstain at P1. Graph showing distribution of BrdU positive neurons in *Pten^T366A/T366A^* and wt mice at P1. **(F)** Example images of BrdU labelling and HOECHST counterstain at P8. Graph showing distribution of BrdU positive neurons in *Pten^T366A/T366A^* and wt mice at P8. Each dot in graphs accounts for one brain section. Statistical analysis with unpaired t-test and two-way ANOVA, Bonferroni post hoc test. For analysis details see **Supplementary Table 2**, Scale bars 100 μm.

Because the Pten behavioural phenotype is established early during development ^65,66^, we also tested whether proliferation of cortical progenitors was affected in *Pten^T366A/T366A^* mice. We performed a BrDU proliferation assay in timed pregnant homozygous *Pten^T366A/T366A^*, heterozygous *Pten^T366A/+^*, and wt mice (Inta et al., 2008). We injected BrdU intraperitoneally in pregnant females at embryonic stages E11.5, E13.5, and E15.5 (**Figure 2, Supplementary Table 2**), and processed brains of offspring for immunohistochemistry of BrdU at P1 and P8 (**Figure 2E, 2F**). Cortical layer formation in the mouse cortex starts at E10.5 with formation of the preplate and projection neurons develop in a tightly controlled temporal and spatial order from E11.5 to E17.5 with L6 and L5, continuing to L4, and L2/3 at stages E15/ E16 ^67,68^. We found BrdU-positive neurons in upper layers 4 to 1. We did not find, however, significant differences in the percentages of BrdU positive neurons across the cortical layers (**Figure 2E, 2F, Supplementary Table 2**). We thus conclude that cortical layers form appropriately in *Pten^T366A/T366A^* mice.

### Soma size in cortical upper layers is affected in *Pten^T366A/T366A^* brains

Previous studies have shown that loss of Pten leads to a developmental state dependent increase in soma size and dendritic arborisation^2,25^. To determine whether Pten T366 affects neuronal growth, we analyzed neuronal soma size in *Pten^T366A/T366A^* cortices at various stages of development (**Figure 3, Supplementary Fig. 2, Supplementary Tables 3, 4**). We first examined soma size of cortical neurons in somatosensory (S1) cortex from *Pten^T366A/T366A^* mice. For this, we injected AAV9-GAG-nls-GFP in postnatal mouse pups at P0, and analyzed GFP positive neurons at P8, P14, P21 and P42 (schematic, **Figure 3A**). We concentrated our analysis on S1 cortex and the granule neurons of the dentate gyrus of the hippocampus. We defined the cortical layers according to the Allen brain atlas, and with HOECHST blue-fluorescent stain. GFP-positive neurons were present in all cortical layers and in the hippocampus (**Figure 3A**). In comparison to wt brains, *Pten^T366A/T366A^* mice showed increased neuronal soma size in pyramidal neurons in L2/3 and in L4 interneurons by P8 and P14 (**Figure 3A, Supplementary Fig. 2A**). In contrast by P21 and P42, L5 pyramidal neurons in *Pten^T366A/T366A^* did not show any differences in size when compared to wt (**Figure 3C, Supplementary Fig. 2B**).

**Figure 3.**
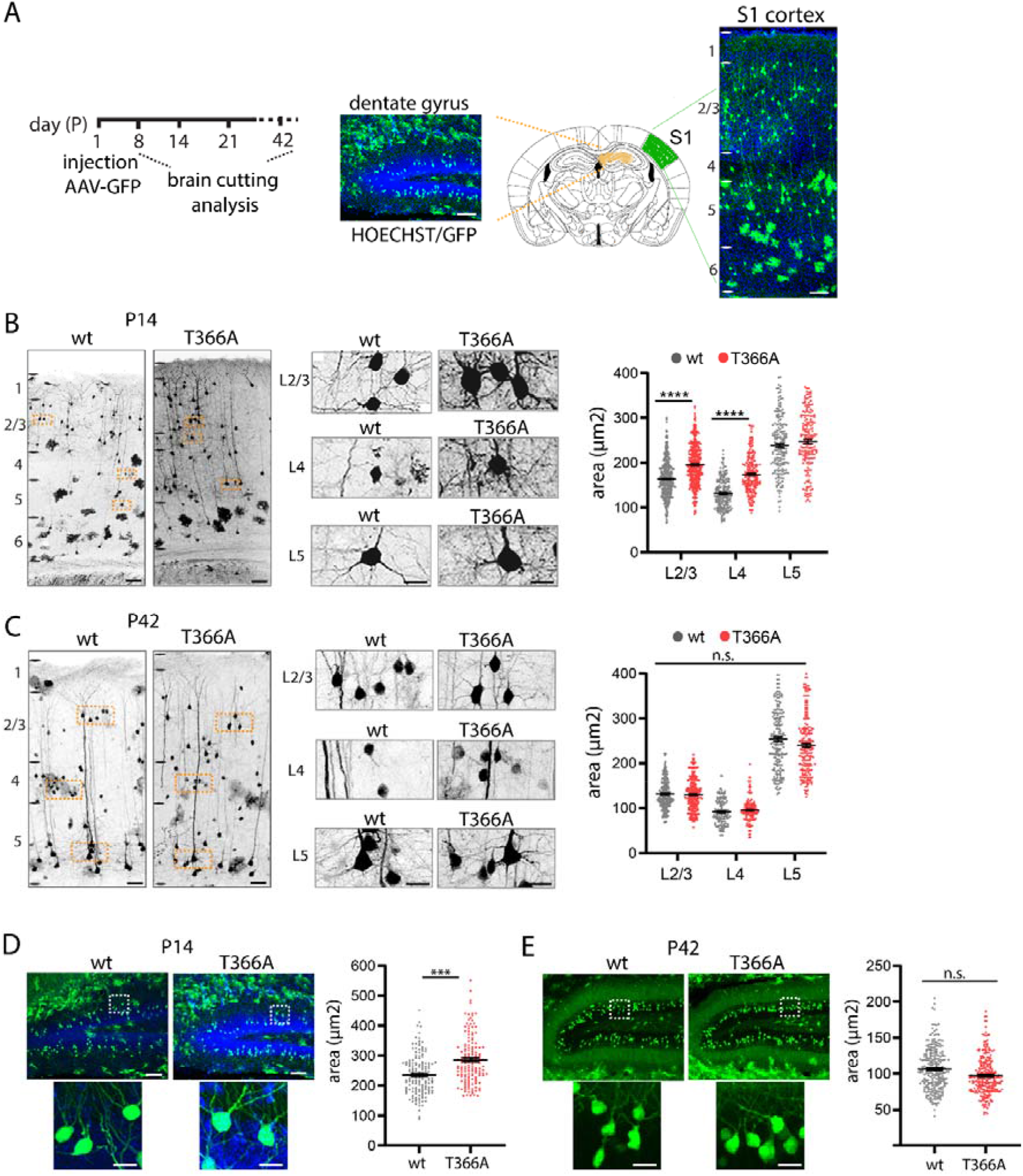
Soma size in *Pten^T366A/T366A^* cortical and dentate gyrus neurons. **(A)** AAV-GFP injection at P0 and expression of GFP in adult S1 cortex and dentate gyrus (P42). Soma size was analyzed at P14 and P42. Staining with HOECHST and the Allen brain mouse reference atlas were used to define cortical layers (also see method section). **(B)** GFP-injected brain slices at P14 for *Pten^T366A/T366A^* and wt mice. Zoom-ins showing somata from pyramidal neurons in L2/3 and L5, and neurons in L4. Graphs at P14 for soma size in L2/3 to L5 *Pten^T366A/T366A^* and wt neurons. **(C)** GFP-injected neurons at stage P42 in *Pten^T366A/T366A^* and wt mice. Graphs at P42 for soma size for L2/3 to L5 *Pten^T366A/T366A^* and wt neurons. **(D)** Analysis of soma size in dentate gyrus neurons at P14 in *Pten^T366A/T366A^* and wt mice. Each dot in graphs accounts for one cell. Data shown as average +/− S.E.M. Data from three brains each genotype, 6 sections per brain. Statistical analysis with one-way ANOVA, ***p<0.001, unpaired t-test for dentate gyrus. Analysis details **Supplementary Table 3**, Scale bars 100 μm, in zoom-ins in B 20 μm, in C and D, E, 50 μm.

We obtained similar results in the dentate gyrus. Labelled pyramidal neurons in dentate gyrus of *Pten^T366A/T366A^* mice were significantly larger than somata in wt mice at P14 (**Figure 3D**), whilst by P42 the soma size was similar in both genotypes (**Figure 3E**). Statistical analysis indicates that soma size in *Pten^T366A/T366A^* mice increased significantly between postnatal stage P8 and P21 with a peak of increase at P14.

Control measurements in primary neuronal cultures plated at E16.5 and analyzed at DIV2-4, 8, and 14 were consistent with these results (**Supplementary Fig. 3A, Supplementary Table 4**). Immunolabeling with MAP2 was performed to label the neurons. The cortical and hippocampal neurons showed no differences in soma size (**Supplementary Fig. 3B-3D**). These results argue for layer- and neuron-specific difference in somata size in *Pten^T366A/T366A^* mice.

### Dendritic arborisation is increased in *Pten^T366A/T366A^* neurons

We next examined whether, in addition to changes in soma size, the dendritic arborization of *Pten^T366A/T366A^* neurons was modified. We analyzed dendritic complexity and dendritic length of pyramidal L2/3 and L4 interneurons by Sholl analysis ^63^ (**Figure 4, Supplementary Fig. 4**). Analysis of dendritic arbors of pyramidal neurons in L2/3 and of interneurons in L4 in S1 cortex revealed an increase in dendritic arborisation in *Pten^T366A/T366A^* brains from P8 onwards.

**Figure 4.**
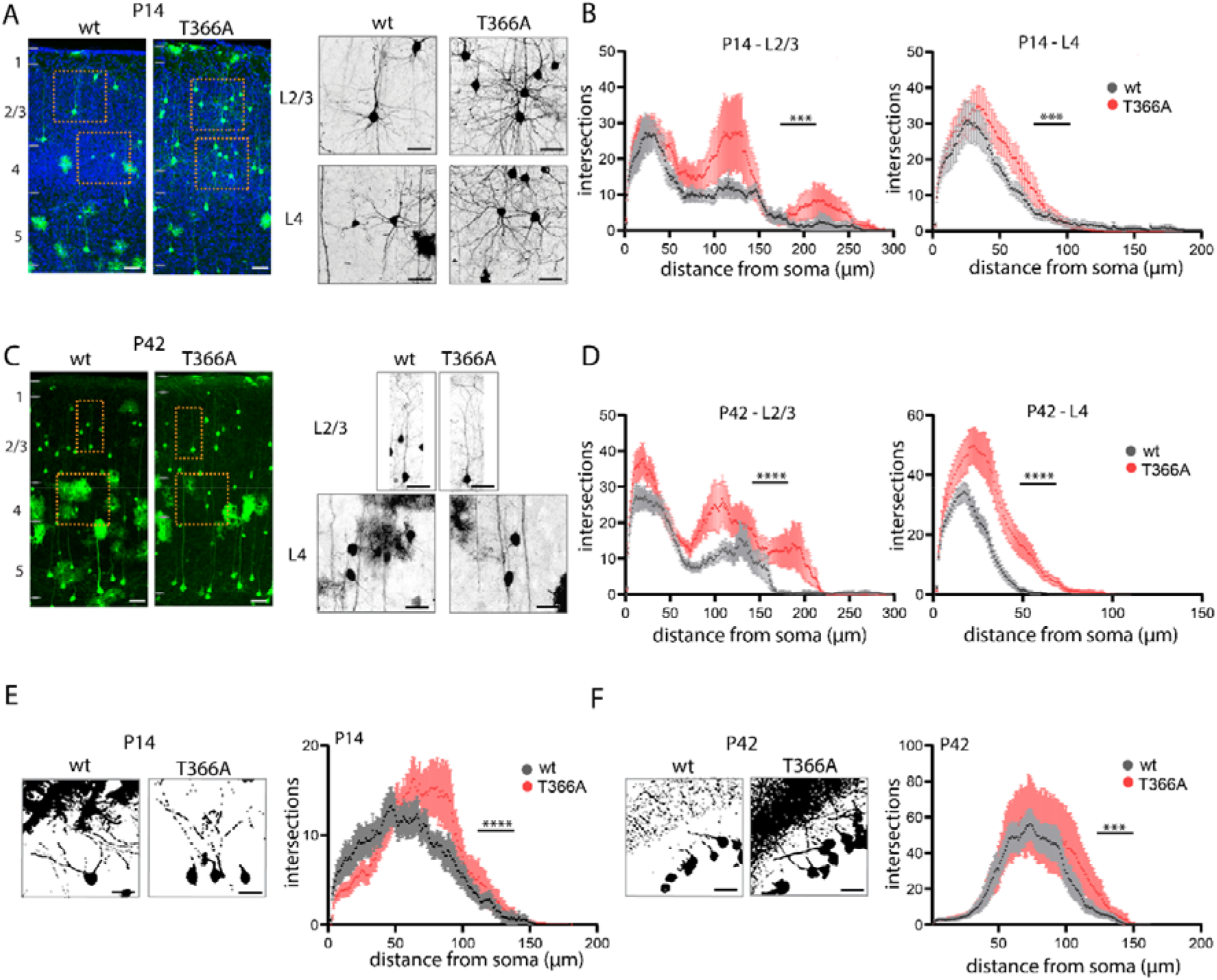
Dendritic arborisation in *Pten^T366A/T366A^* cortical and dentate gyrus neurons. **(A, B)** Dendritic arborization for L2/3 pyramidal neurons and L4 neurons at P14. Zoom-in for L2/3 pyramidal neurons and L4 interneurons. Graphs showing quantification with Sholl analysis. (**C, D**) Dendritic arborisation for L2/3 and L4 neurons at P14 and P42. (**E**) Zoom-in for L2/3 pyramidal neurons and L4 neurons. Graphs showing quantification with Sholl analysis. **(E, F)** Dendritic arborization in hippocampal dentate gyrus neurons at P14 and P42 with quantification in graphs. Data shown as average +/− S.E.M. Statistical analysis with twoway ANOVA, Bonferroni post hoc test, ***p<0.001. Scale bars 100 μm, 50 μm in zoom-ins.

At P14, the dendritic arborization of *Pten^T366A/T366A^* L2/3 pyramidal neurons and *Pten^T366A/T366A^* L4 interneurons were significantly increased (**Figure 4A, 4B**). In *Pten^T366A/T366A^* L2/3 neurons the extent of arborisation differed from wt in the region of the oblique dendrites (70 μm to 150 μm from the soma) and in the region of the apical dendrites (200 μm to 250 μm from the soma). *Pten^T366A/T366A^* L4 neurons showed a significant increase in the number of proximal branches at 30 μm to 70 μm from the soma. At P42, dendritic arborisation in *Pten^T366A/T366A^* neurons was significantly increased in L2/3 and L4 (**Figures 4C, 4D**). In *Pten^T366A/T366A^* L2/3 neurons the extent of arborisation differed from wt in the region of the oblique dendrites and in the region of the apical dendrites similarly to P14. *Pten^T366A/T366A^* L4 neurons showed a significant increase in the number of proximal branches at 10 μm to 50 μm from the soma. In contrast to the results for the soma size, the differences in dendritic arborization persisted from P8 to P42. In addition, the total dendritic lengths were significantly increased in *Pten^T366A/T366A^* L2/3 pyramidal neurons and L4 interneurons at P14. However, no changes were observed at P42 (**Supplementary Fig. 4A-4D, Supplementary Table 5**).

We performed a similar analysis of hippocampal dentate gyrus granule neurons (**Figure 4E, 4F**), and found the same effect as with cortical neurons. Dendritic arbors in *Pten^T366A/T366A^* mice were larger at both stages P14 and P42 compared to wt. Sholl analysis of all order of branches for hippocampal dentate gyrus granule neurons at P14 showed particularly increased arborisation of *Pten^T366A/T366A^* neurons from 50 μm to 100 μm distance from the soma. By comparison, wt dentate gyrus dendrites were normally distributed with a peak at 50 μm from soma (**Figure 4E**). At P42, dendritic arbors in *Pten^T366A/T366A^* neurons showed more complexity between 100 μm and 140 μm from the soma compared to wt (**Figure 4F**). The average length of all dendrites was increased for *Pten^T366A/T366A^* and wt neurons at P14 in L2/3 dendrites, and at P42 for L4 dendrites (**Supplementary Fig. 4E, 4F, Supplementary Table 5**). Taken together, these results demonstrate that cortical morphology is affected in *Pten^T366A/T366A^* mice.

### Disruption of the thalamo-cortical / cortico-cortical balance in *Pten^T366A/T366A^* mice

Next we examined whether morphological differences translate into changes in neuronal connectivity. Earlier work using post-mortem tissue ^69,70^, and magnetic resonance tomography connectivity mapping ^71,72^ suggests that cortical connectivity might be affected in ASD. In humans, Pten germline variations lead to imbalances of cortical connections ^66,73^. Similarly, in a Pten mouse model for autism (*Pten^+/−^*), connectivity between prefrontal cortex and amygdala is disturbed ^16,74^.

To characterize whether the dendritic arborization phenotype in *Pten^T366A/T366A^* cortical neurons is reflected in cortical connectivity, we performed retrograde rabies virus tracing. Retrograde rabies virus tracing has been widely used to track presynaptic input to a defined cortical area and class of neurons, i.e. interneurons or pyramidal cells ^41,75,76^. In the Pten mice, the number of potential changes is vast, and the class of laminar or cell type specific mice lines are difficult to establish. One hypothesis related to ASD and Pten mice is that the balance between cortico-cortical and thalamo-cortical connectivity is modified ^72,77,78^. To examine this, we combined anterograde transsynaptic tracing with retrograde monosynaptic rabies tracing ^79^. We used bulk injections of a trans-synaptic virus -- AAV1-Cre into ventrobasal thalamus. This step puts Cre into the cortical neurons receiving input from higher order thalamus (POM) and the primary somatosensory thalamus (VPm). Next we used Cre-dependent rabies in cortex, to examine how cortical connectivity related to this class of neurons connected to thalamus changed (**Figure 5, Supplementary Fig. 5, Supplementary Table 6**).

**Figure 5.**
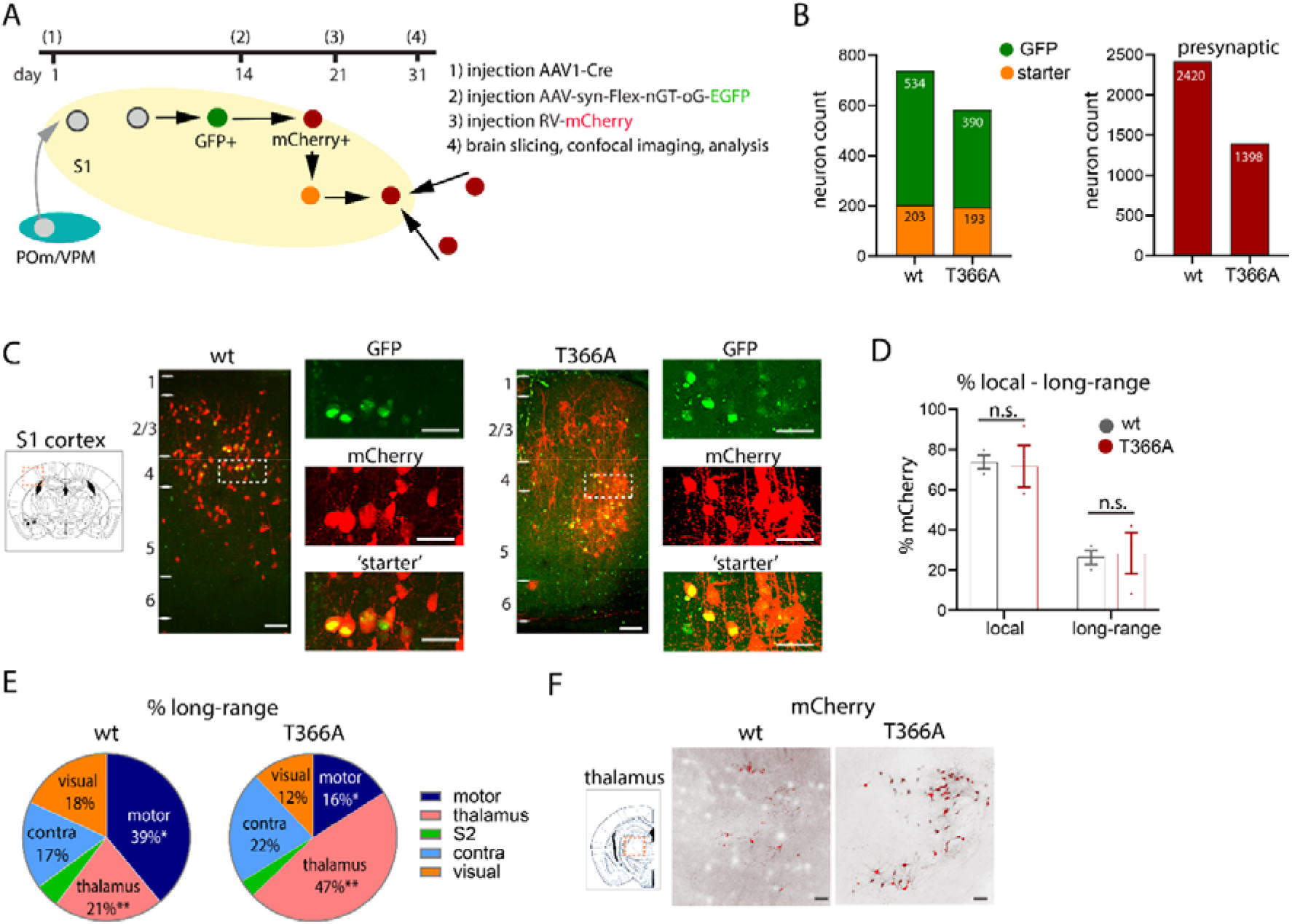
Presynaptic input to S1 cortex in *Pten^T366A/T366A^* mice. **(A)** Schematic showing injection procedure. **(B)** Graph showing number of GFP-positive and starter neurons (*left*), and presynaptic neurons (*right*). (**C**) Example images showing expression of AAV-FleX-EGFP and presynaptic RV-mCherry neurons in wt and *Pten^T366A/T366A^* S1 cortex. Zoom-ins showing starter neurons in yellow. (**D**) Graph showing the percentage of presynaptic neurons local in S1, and long-range neurons in *Pten^T366A/T366A^* and wt brains. **(E)** Pie charts showing percentages of long-range presynaptic input to S1 in *Pten^T366A/T366A^* and wt brains. **(F)** Example images of presynaptic neurons in thalamus in *Pten^T366A/T366A^* and wt brains. Statistical analysis with unpaired t-test, *p<0.05. Analysis details in **Supplementary Table 6,** scale bars 100 μm, 50 μm in zoom-ins.

We first injected an anterograde AAV1-Cre in the ventrobasal thalamus. The axons of these neurons target L5a and L1 (POm thalamus) and L4 (VPm thalamus) in S1 cortex ^80,81^. In this configuration, the AAV is expressed in axon terminals in L1, L2/3, L4, L5 and L6a of S1 cortex and AAV1-Cre is expressed in postsynaptic targets of these axons in S1 cortex. A second injection in S1 cortex was then used to target Cre-positive neurons in S1 with a Flex-switch virus (AAV1-Syn-Flex-nGToG-WPRE3) ^82^. Finally, a third injection using RABV-SADB19dG-mCherry was used to trace the previously infected neurons. We analyzed the expression of presynaptic neurons 10 days after the last injection (**Figure 5A, Supplementary Fig. 5A**).

We analyzed every second section of three brains for each genotype. We counted 203 starter neurons out of 534 GFP positive neurons for wt, and 193 starter neurons out of 390 GFP positive neurons for *Pten^T366A/T366A^* brains (**Figure 5B, *left***). A total of 2420 presynaptic neurons were found in the wt brains, and a total of 1398 presynaptic neurons in the *Pten^T366A/T366A^* brains (**Figure 5C, *right***). Example images of S1 cortex of wt and *Pten^T366A/T366A^* brains show specific labelling of GFP and mCherry expressing (yellow) ‘starter’ neurons in L2/3, L4, and L5 in S1 cortex confirming that the injection in the thalamus successfully labelled Cre-positive cells (**Figure 5C**).

Immunolabeling for Cre-positive neurons in S1 cortex showed that the AAV1-Cre functioned transsynaptically, as there were Cre-positive neurons in S1 in both wt and *Pten^T366A/T366A^* mice (**Supplementary Fig. 5B**). The number and distribution of Cre-positive neurons in cortex in wt and *Pten^T366A/T366A^* mice was similar. Immunolabeling for Cre-positive neurons in VPm/POm thalamus confirmed the presence of label at the ventrobasal injection site of the AAV1-Cre virus (**Supplementary Fig. 5C**).

The cortical connectivity of these neurons receiving input from thalamus was modified in *Pten^T366A/T366A^* brains. In wt mice, local presynaptic axonal input to the starter cells constituted 74% of the total input to these cells, while in *Pten^T366A/T366A^* brains the local input accounted for 72% of the total input to the starter neurons. The long-range presynaptic input to S1 cortex from other cortical areas was 26% in wt compared to 28% in *Pten^T366A/T366A^* brains (**Figure 5D**). Thus, the balance between local and long-range cortical connectivity was unchanged for the cortical neurons synaptically connected to the cortico-thalamic neurons.

However, there was a fundamental shift in the long-range connectivity of these cortical neurons in the *Pten^T366A/T366A^* brains: Dissection of the long-range input showed that in both genotypes, the bulk of the input arose from visual cortices (23% wt, 11% *Pten^T366A/T366A^*); S2 cortex (5% wt, 3% in *Pten^T366A/T366A^*) and contralateral cortex accounted for 17% wt, and 22% in *Pten^T366A/T366A^.* Input to motor cortices (39% wt, 16% *Pten^T366A/T366A^*) and thalamic areas (21% wt, 47% *Pten^T366A/T366A^*) were significantly different in *Pten^T366A/T366A^* brains (**Figure 5E, 5F**).

The significant difference in the input from thalamus to S1 cortex and from motor cortices to S1 cortex between the wt and *Pten^T366A/T366A^* mice suggest that aspects of the cortical and the thalamo-cortical circuit organization have been modified in *Pten^T366A/T366A^* mice. As a control, we also performed commonly used rabies virus tracing in *Pten^T366A/T366A^* and wt mice (data not shown) without first using the trans-synaptic AAV1-Cre in thalamus. In these experiments we did not observe any significant difference in local versus long-range projection to S1 cortex, and there were no differences in the proportion of thalamic neurons presynaptic to S1 neurons, suggesting that the difference in cortical circuit organization in *Pten^T366A/T366A^* brains arises from some modification in aspects of cortical and thalamo-cortical connectivity. In conclusion, the connectivity tracing results suggest that presynaptic input from motor cortex and thalamic input to S1 cortex differ in *Pten^T366A/T366A^* brains and lead to an imbalance in long-range input.

## Discussion

We present results from a newly generated mutant mouse line carrying a Pten mutation at the phosphorylation site threonine 366. Knock-in *Pten^T366A/T366A^* mice are viable and born in a Mendelian ratio, and therefore provide an excellent model *in vivo* for investigation of this phosphorylation site. *Pten^T366A/T366A^* mice show abnormal cognitive aspects of behaviour. These neurological deficits presumably arise from developmentally regulated changes in soma size and dendritic arbors of cortical neurons, and by consequence long-term changes in connectivity.

### Behaviour in *Pten^T366A/T366A^* mice

Previous work has shown that *Pten Nse-Cre* mutant mice have deficits in social activity, reaction to sensory stimuli and have increased anxiety-like behaviours^2,15,83^. Also in *Pten^+/−^* mice, social behaviour was impaired and associated with increased brain size, and considered an ASD-related characteristic^17,18^. In *Pten^T366A/T366A^* mice, exploratory behaviour in the open field test and motor behaviour in the hanging wire and rotarod tests were similar to that observed in wt littermates, suggesting that general motor function is intact in these mice. However, the deficits in spatial working memory (Y maze), spatial navigation (Barnes maze and Morris water maze), contextual fear conditioning and vibrissae-stimulated reflexes in *Pten^T366A/T366A^* mice support the idea that sensory integration with hippocampal and entorhinal cortical circuitries involved in spatial cognition ^84^ is impaired in *Pten^T366A/T366A^* mice.

### Cortical morphology in *Pten^T366A/T366A^* mice

Earlier work has shown that *Pten* mutations affect neuron size ^2,15^. A compartmental increase in pyramidal neuron dendrites has also been demonstrated in *Pten^+/−^* mice ^16,25^. In line with these earlier studies, the dendritic arbor of cortical pyramidal neurons in S1 cortex is enlarged in *Pten^T366A/T366A^* mice even though cortical proliferation and layer formation were not affected. Whether the dendritic phenotype of L2/3 pyramidal neurons in *Pten^T366A/T366A^* mice influences the activity of apical tufts in cortical L1, and whether the activity of L5 apical tuft dendrites in L1 is modified in a perceptual task is unknown ^85^.

In contrast to the increased dendritic arborization of pyramidal neurons, the effect of phosphorylation at Pten T366 on soma size is developmentally regulated. It is restricted to neurons in upper L2/3 and to L4 in S1 cortex and transiently expressed from P8 to P14. Soma size reverts to normal levels from P21 onwards. The molecular mechanisms underlying the effects of phosphorylation at Pten T366 on soma size in these neurons is not clear.

Several human *PTEN* ASD mutations affect protein stability and activity ^5^. Interestingly, an ASD patient has been identified, who shows a decrease in phosphorylation at Pten T366 ^86^, thus mimicking the PTEN T366A phenotype. Testing the phosphorylation site at Pten T366 for protein stability and activity could provide an entry point for understanding the effect of phosphorylation at Pten T366 on soma size or neuron size in general. Further, it could help in understanding the involvement of at Pten T366 phosphorylation in the context of ASD.

### Cortical connectivity in *Pten^T366A/T366A^* mice

In line with the altered morphology of pyramidal neurons in *Pten^T366A/T366A^* mice, connectivity tracing with rabies virus showed an imbalance of long-range cortical input to S1 cortex in *Pten^T366A/T366A^* mice. Neurons in *Pten^T366A/T366A^* mice had a smaller number of presynaptic inputs from motor cortices. Along with this modification in the cortical connectivity, the thalamo-cortical pathway was disrupted in *Pten^T366A/T366A^* mice. Neurons in cortex that received thalamic were better connected in *Pten^T366A/T366A^* mice than in wt mice. In this context the behavioural changes in *Pten^T366A/T366A^* mice could arise from changes in long-range connectivity, which could result in alterations in sensory processing. Our results suggest that phosphorylation at Pten T366 is implicated in interactions between feedforward and feedback connectivity. Furthermore, imbalance of cortical connections have been described in ASD patients^72,77,78^. To dissect the circuit properties in *Pten^T366A/T366A^* mice that underlie Pten T366 function, a functional analysis of dendritic activity by two-photon Ca^2+^ imaging or of long-range input to somatosensory cortex in *Pten^T366A/T366A^* mice (by optogenetic approaches) would prove useful ^39,87^.

### Conclusion

Our analyses of the *Pten^T366A/T366A^* mice show that phosphorylation of Pten at T366 impacts on cortical cell morphology and connectivity, which may account for changes in behavioural characteristics. Our analysis of T366 contributes to the understanding of Pten function in sensory processing with a potential link to ASD.

## Acknowledgements

We thank Dr. Thorsten Trimbuch and the Viral Core facility for their support and the generation of the viruses used in this study. We thank Dr. Jan Schmoranzer and the AMBIO facility for their support and the use of the microscope facility. We like to thank Zara Khan for helping with the analysis of dendrite arborisation. Finally, we thank Kristin Lehmann, Kerstin Schlawe and Beate Dietmar for excellent technical assistance.

## Supplementary material

Supplementary material is available at *Brain* online.

## Author contributions

J.L., J.A.B., R.S., F.F. and B.J.E. planned and designed the experiments and wrote the manuscript. J.L., J.A.B., R.R. and A.J.R. performed the experiments. J.L., J.B., and J.A.R. analyzed the data. All authors approved the final version of the paper.

## Abbreviations

ASD: Autism spectrum disorders
DIV: days *in vitro*
E: embryonic day
L: layer
PFA: paraformaldehyde
P: postnatal day
S1: primary somatosensory cortex

---

**Supplementary Figure 1 Protein levels in *Pten^T366A/T366A^* brains.** Protein levels of pAkt, Pten, pS6 and pGSK3b in forebrain lysates from P10 brains of *Pten^T366A/T366A^* and wt mice. Molecular weight protein ladder is in kilodaltons.

**Supplementary Figure 2 Soma size in *Pten^T366A/T366A^* cortical neurons at P8 and P21. (A)** Images showing GFP expression in S1 cortex at P8. Zoom-ins showing somata from pyramidal neurons in L2/3 and L5, and neurons in L4. Graphs at P8 for soma size in L2/3 to L5 neurons. **(B)** Images showing GFP expression in S1 cortex at P21. Graph showing quantitative analysis for somata size. Each dot in graphs accounts for one cell. Data shown as average +/− S.E.M. Statistical analysis with one-way ANOVA, **p<0.01. For analysis details see **Table S2**, Scale bars 100 μm, 50 μm in zoom-ins.

**Supplementary Figure 3 Soma size in *Pten^T366A/T366A^* primary cortical and hippocampal neurons. (A)** Soma size was analyzed in primary cell culture from E16.5 hippocampal and cortical neurons with MAP2 immunostaining. (**B-E**) Example images from cortical neurons (**B**) and hippocampal neurons (**D)** at DIV 2-3, DIV 8 and DIV 14. (**D**) Graphs showing analysis for cortical (**C**) and hippocampal (**E**) neurons. Each dot in graphs accounts for one cell. Data shown as average +/− S.E.M. Two pregnant mice each genotype and cell type, cortices/ hippocampi of 5-8 embryos were pooled. Statistical analysis with one-way ANOVA, details **Supplementary Table 4**, Scale bars 50 μm.

**Supplementary Figure 4 Dendritic lengths of cortical and dentate gyrus neurons in *Pten^T366A/T366A^* and wt mice. (A, B)** Images showing reconstruction of dendritic length of L2/3 pyramidal neurons (**A**), and of L4 neurons (**B**) in *Pten^T366A/T366A^* and wt mice at P14 and at P42 (**C, D**) in *Pten^T366A/T366A^* and wt mice. (**E, F**) Images of reconstruction of dentate gyrus at P14 (**E**) and P42 (**F**) in wt and *Pten^T366A/T366A^* mice. Data from three brains each genotype. Statistical analysis was performed with unpaired t-test, ***p<0.001. Analysis details in **Supplementary Table 5**.

**Supplementary Figure 5 Injection sites and Cre expression in wt and in *Pten^T366A/T366A^* mice. (A, B)** Images showing injection sites in S1 cortex in wt (*top*) and *Pten^T366A/T366A^* (*below*) mice. (**C, D**) Example images showing Cre expression and presynaptic mCherry positive neurons in S1 cortex and in thalamus. Scale bars 500 μm in A, 500 μm, in zoom-ins 200 μm; in B, C, 100 μm, 50 μm in zoom-in.

## References

1. Cohen MM. The AKT genes and their roles in various disorders. Am J Med Genet Part A. 2013;161:2931–2937. doi:10.1002/ajmg.a.36101

2. Kwon CH, Luikart BW, Powell CM, et al. Pten Regulates Neuronal Arborization and Social Interaction in Mice. Neuron. 2006;50(3):377–388. doi:10.1016/j.neuron.2006.03.023

3. Lv JW, Cheng TL, Qiu ZL, Zhou WH. Role of the PTEN signaling pathway in autism spectrum disorder. Neurosci Bull. 2013;29(6):773–778. doi:10.1007/s12264-013-1382-3

4. Rademacher S, Eickholt BJ. PTEN in Autism and Neurodevelopmental Disorders. Cold Spring Harb Perspect Med. August 2019. doi:10.1101/cshperspect.a036780

5. Spinelli L, Black FM, Berg JN, Eickholt BJ, Leslie NR. Functionally distinct groups of inherited PTEN mutations in autism and tumour syndromes. 2015:128–134. doi:10.1136/jmedgenet-2014-102803

6. Rodríguez-Escudero I, Oliver MD, Andrés-Pons A, Molina M, Cid VJ, Pulido R. A comprehensive functional analysis of PTEN mutations: implications in tumor- and autism-related syndromes. Hum Mol Genet. 2011;20(21):4132–4142. doi:10.1093/hmg/ddr337

7. Carter MT, Scherer SW. Autism spectrum disorder in the genetics clinic: a review. Clin Genet. 2013;83(5):399–407. doi:10.1111/cge.12101

8. Fernandez M, Mollinedo-Gajate I, Penagarikano O. Neural Circuits for Social Cognition: Implications for Autism. Neuroscience. 2018;370:148–162. doi:10.1016/j.neuroscience.2017.07.013

9. Balci TB, Davila J, Lewis D, et al. Broad spectrum of neuropsychiatric phenotypes associated with white matter disease in PTEN hamartoma tumor syndrome. Am J Med Genet Part B, Neuropsychiatr Genet Off Publ Int Soc Psychiatr Genet. 2018;177(1):101–109. doi:10.1002/ajmg.b.32610

10. Vanderver A, Tonduti D, Kahn I, et al. Characteristic brain magnetic resonance imaging pattern in patients with macrocephaly and PTEN mutations. Am J Med Genet A. 2014;164A(3):627–633. doi:10.1002/ajmg.a.36309

11. Hansen-Kiss E, Beinkampen S, Adler B, et al. A retrospective chart review of the features of PTEN hamartoma tumour syndrome in children. J Med Genet. 2017;54(7):471–478. doi:10.1136/jmedgenet-2016-104484

12. Busch RM, Srivastava S, Hogue O, et al. Neurobehavioral phenotype of autism spectrum disorder associated with germline heterozygous mutations in PTEN. Transl Psychiatry. 2019;9(1):253. doi:10.1038/s41398-019-0588-1

13. Clipperton-Allen AE, Cohen OS, Aceti M, et al. Pten haploinsufficiency disrupts scaling across brain areas during development in mice. Transl Psychiatry. 2019;9(1):329. doi:10.1038/s41398-019-0656-6

14. Frazier TW, Embacher R, Tilot AK, Koenig K, Mester J, Eng C. Molecular and phenotypic abnormalities in individuals with germline heterozygous PTEN mutations and autism. Mol Psychiatry. 2015;20(9):1132–1138. doi:10.1038/mp.2014.125

15. Amiri a., Cho W, Zhou J, et al. Pten Deletion in Adult Hippocampal Neural Stem/Progenitor Cells Causes Cellular Abnormalities and Alters Neurogenesis. J Neurosci. 2012;32(17):5880–5890. doi:10.1523/JNEUROSCI.5462-11.2012

16. Huang W-C, Chen Y, Page DT, et al. Hyperconnectivity of prefrontal cortex to amygdala projections in a mouse model of macrocephaly/autism syndrome. Nat Commun. 2016;7:13421. doi:10.1038/ncomms13421

17. Page DT, Kuti OJ, Prestia C, Sur M. Haploinsufficiency for Pten and Serotonin transporter cooperatively influences brain size and social behavior. Proc Natl Acad Sci U S A. 2009;106(6):1989–1994. doi:10.1073/pnas.0804428106

18. Clipperton-Allen AE, Page DT. Pten haploinsufficient mice show broad brain overgrowth but selective impairments in autism-relevant behavioral tests. Hum Mol Genet. 2014;23(13):3490–3505. doi:10.1093/hmg/ddu057

19. Vanhaesebroeck B, Stephens L, Hawkins P. PI3K signalling: the path to discovery and understanding. Nat Rev Mol Cell Biol. 2012;13(3):195–203. doi:10.1038/nrm3290

20. Kwon CH, Zhu X, Zhang J, et al. Pten regulates neuronal soma size: a mouse model of Lhermitte-Duclos disease. Nat Genet. 2001;29(4):404–411. doi:10.1038/ng781

21. van Diepen MT, Eickholt BJ. Function of PTEN during the formation and maintenance of neuronal circuits in the brain. Dev Neurosci. 2008;30(1-3):59–64. doi:10.1159/000109852

22. Jaworski J, Spangler S, Seeburg DP, Hoogenraad CC, Sheng M. Control of dendritic arborization by the phosphoinositide-3’-kinase-Akt-mammalian target of rapamycin pathway. J Neurosci. 2005;25(49):11300–11312. doi:10.1523/JNEUROSCI.2270-05.2005

23. van Diepen MT, Parsons M, Downes CP, Leslie NR, Hindges R, Eickholt BJ. MyosinV controls PTEN function and neuronal cell size. Nat Cell Biol. 2009;11(10):1191–1196. doi:10.1038/ncb1961

24. Kwon C-H, Zhu X, Zhang J, Baker SJ. mTor is required for hypertrophy of Pten-deficient neuronal soma in vivo. Proc Natl Acad Sci U S A. 2003;100(22):12923–12928. doi:10.1073/pnas.2132711100

25. Chow DK, Groszer M, Pribadi M, et al. Laminar and compartmental regulation of dendritic growth in mature cortex. Nat Neurosci. 2009;12(2):116–118. doi:10.1038/nn.2255

26. Garcia-Junco-Clemente P, Chow DK, Tring E, Lazaro MT, Trachtenberg JT, Golshani P. Overexpression of calcium-activated potassium channels underlies cortical dysfunction in a model of PTEN-associated autism. Proc Natl Acad Sci U S A. 2013;110:18297–18302. doi:10.1073/pnas.1309207110

27. Groszer M, Erickson R, Scripture-Adams DD, et al. Negative regulation of neural stem/progenitor cell proliferation by the Pten tumor suppressor gene in vivo. Science. 2001;294(5549):2186–2189. doi:10.1126/science.1065518

28. Planchon SM, Waite K a, Eng C. The nuclear affairs of PTEN. J Cell Sci. 2008;121(Pt 3):249–253. doi:10.1242/jcs.022459

29. Fricano-Kugler CJ, Getz SA, Williams MR, et al. Nuclear Excluded Autism-Associated Phosphatase and Tensin Homolog Mutations Dysregulate Neuronal Growth. Biol Psychiatry. 2018;84(4):265–277. doi:10.1016/j.biopsych.2017.11.025

30. Mingo J, Rodríguez-Escudero I, Luna S, et al. A pathogenic role for germline PTEN variants which accumulate into the nucleus. Eur J Hum Genet. 2018;26(8):1180–1187. doi:10.1038/s41431-018-0155-x

31. Takumi T, Tamada K, Hatanaka F, Nakai N, Bolton PF. Behavioral neuroscience of autism. Neurosci Biobehav Rev. 2020;110:60–76. doi:10.1016/j.neubiorev.2019.04.012

32. Winden KD, Ebrahimi-Fakhari D, Sahin M. Abnormal mTOR Activation in Autism. Annu Rev Neurosci. 2018;41(1):1–23. doi: 10.1146/annurev-neuro-080317-061747

33. Xu D, Yao Y, Jiang X, Lu L, Dai W. Regulation of PTEN stability and activity by Plk3. J Biol Chem. 2010;285(51):39935–39942. doi:10.1074/jbc.M110.166462

34. Al-khouri AM, Ma Y, Williams S, Mustelin T. Cooperative Phosphorylation of the Tumor Suppressor Phosphatase and Tensin Homologue (PTEN) by Casein Kinases and Glycogen Synthase Kinase 3 □ *. 2005;280(42):35195–35202. doi:10.1074/jbc.M503045200

35. Tibarewal P, Zilidis G, Spinelli L, et al. PTEN Protein Phosphatase Activity Correlates with Control of Gene Expression and Invasion, a Tumor-Suppressing Phenotype, But Not with AKT Activity. Sci Signal. 2012;5(213):ra18–ra18. doi:10.1126/scisignal.2002138

36. Maccario H, Perera NM, Davidson L, Downes CP, Leslie NR. PTEN is destabilized by phosphorylation on Thr366. Biochem J. 2007;405:439–444. doi:10.1042/BJ20061837

37. Okumura K, Zhao M, Depinho R a, Furnari FB, Cavenee WK. Cellular transformation by the MSP58 oncogene is inhibited by its physical interaction with the PTEN tumor suppressor. Proc Natl Acad Sci U S A. 2005;102(8):2703–2706. doi:10.1073/pnas.0409370102

38. Schrötter S, Leondaritis G, Eickholt BJ. Capillary Isoelectric Focusing of Akt Isoforms Identifies Highly Dynamic Phosphorylation in Neuronal Cells and Brain. 2016;291(19):10239–10251. doi:10.1074/jbc.M115.700138

39. Zolnik TA, Ledderose J, Toumazou M, et al. Layer 6b Is Driven by Intracortical Long-Range Projection Neurons. Cell Rep. 2020;30(10):3492–3505.e5. doi:10.1016/j.celrep.2020.02.044

40. Sun Y, Nguyen AQ, Nguyen JP, et al. Cell-type-specific circuit connectivity of hippocampal CA1 revealed through Cre-dependent rabies tracing. Cell Rep. 2014;7(1):269–280. doi:10.1016/j.celrep.2014.02.030

41. Kim EJ, Jacobs MW, Ito-Cole T, Callaway EM. Improved Monosynaptic Neural Circuit Tracing Using Engineered Rabies Virus Glycoproteins. Cell Rep. 2016;15(4):692–699. doi:10.1016/j.celrep.2016.03.067

42. Osakada F, Mori T, Cetin AH, Marshel JH, Virgen B, Callaway EM. New rabies virus variants for monitoring and manipulating activity and gene expression in defined neural circuits. Neuron. 2011;71(4):617–631. doi:10.1016/j.neuron.2011.07.005

43. Pilpel N, Landeck N, Klugmann M, Seeburg PH, Schwarz MK. Rapid, reproducible transduction of select forebrain regions by targeted recombinant virus injection into the neonatal mouse brain. J Neurosci Methods. 2009;182:55–63. doi:10.1016/j.jneumeth.2009.05.020

44. Inta D, Alfonso J, von Engelhardt J, et al. Neurogenesis and widespread forebrain migration of distinct GABAergic neurons from the postnatal subventricular zone. Proc Natl Acad Sci U S A. 2008;105(52):20994–20999. doi:10.1073/pnas.0807059105

45. Lalonde R. The neurobiological basis of spontaneous alternation. Neurosci Biobehav Rev. 2002;26(1):91–104. doi:10.1016/s0149-7634(01)00041-0

46. Arenkiel BR, Ehlers MD. Molecular genetics and imaging technologies for circuit-based neuroanatomy. Nature. 2009;461(7266):900–907. doi:10.1038/nature08536

47. Holcomb LA, Gordon MN, Jantzen P, Hsiao K, Duff K, Morgan D. Behavioral changes in transgenic mice expressing both amyloid precursor protein and presenilin-1 mutations: lack of association with amyloid deposits. Behav Genet. 1999;29(3):177–185. doi:10.1023/a:1021691918517

48. Pennanen L, Wolfer DP, Nitsch RM, Götz J. Impaired spatial reference memory and increased exploratory behavior in P301L tau transgenic mice. Genes Brain Behav. 2006;5(5):369–379. doi:10.1111/j.1601-183X.2005.00165.x

49. Crawley JN. Social Behavior Tests for Mice. What’s Wrong With My Mouse Behav Phenotyping Transgenic Knockout Mice. 2007:65–70.

50. Freitag S, Schachner M, Morellini F. Behavioral alterations in mice deficient for the extracellular matrix glycoprotein tenascin-R. Behav Brain Res. 2003;145(1-2):189–207. doi:10.1016/s0166-4328(03)00109-8

51. Carter RJ, Morton J, Dunnett SB. Motor coordination and balance in rodents. Curr Protoc Neurosci. 2001;Chapter 8:Unit 8.12. doi:10.1002/0471142301.ns0812s15

52. Crawley JN. Behavioral phenotyping of transgenic and knockout mice: experimental design and evaluation of general health, sensory functions, motor abilities, and specific behavioral tests. Brain Res. 1999;835(1):18–26. doi:10.1016/s0006-8993(98)01258-x

53. Deuis JR, Dvorakova LS, Vetter I. Methods Used to Evaluate Pain Behaviors in Rodents. Front Mol Neurosci. 2017;10:284. doi:10.3389/fnmol.2017.00284

54. Lekic T, Manaenko A, Rolland W, et al. Rodent neonatal germinal matrix hemorrhage mimics the human brain injury, neurological consequences, and post-hemorrhagic hydrocephalus. Exp Neurol. 2012;236(1):69–78. doi:10.1016/j.expneurol.2012.04.003

55. Barnes CA. Memory deficits associated with senescence: a neurophysiological and behavioral study in the rat. J Comp Physiol Psychol. 1979;93(1):74–104. doi:10.1037/h0077579

56. Bach ME, Hawkins RD, Osman M, Kandel ER, Mayford M. Impairment of spatial but not contextual memory in CaMKII mutant mice with a selective loss of hippocampal LTP in the range of the theta frequency. Cell. 1995;81(6):905–915. doi:10.1016/0092-8674(95)90010-1

57. Holmes A, Wrenn CC, Harris AP, Thayer KE, Crawley JN. Behavioral profiles of inbred strains on novel olfactory, spatial and emotional tests for reference memory in mice. Genes Brain Behav. 2002;1(1):55–69. doi:10.1046/j.1601-1848.2001.00005.x

58. Paylor R, Zhao Y, Libbey M, Westphal H, Crawley JN. Learning impairments and motor dysfunctions in adult Lhx5-deficient mice displaying hippocampal disorganization. Physiol Behav. 2001;73(5):781–792. doi:10.1016/s0031-9384(01)00515-7

59. Puzzo D, Lee L, Palmeri A, Calabrese G, Arancio O. Behavioral assays with mouse models of Alzheimer’s disease: practical considerations and guidelines. Biochem Pharmacol. 2014;88(4):450–467. doi:10.1016/j.bcp.2014.01.011

60. Maren S. Neurobiology of Pavlovian fear conditioning. Annu Rev Neurosci. 2001;24:897–931. doi:10.1146/annurev.neuro.24.1.897

61. Rudy JW, Huff NC, Matus-Amat P. Understanding contextual fear conditioning: insights from a two-process model. Neurosci Biobehav Rev. 2004;28(7):675–685. doi:10.1016/j.neubiorev.2004.09.004

62. Kenney JW, Gould TJ. Nicotine enhances context learning but not context-shock associative learning. Behav Neurosci. 2008;122(5):1158–1165. doi:10.1037/a0012807

63. Uylings HBM, van Pelt J. Measures for quantifying dendritic arborizations. Network. 2002;13(3):397–414.

64. Tilot AK, Gaugler MK, Yu Q, et al. Germline disruption of Pten localization causes enhanced sex-dependent social motivation and increased glial production. Hum Mol Genet. 2014;23(12):3212–3227. doi:10.1093/hmg/ddu031

65. Hobert JA, Embacher R, Mester JL, Frazier TW 2nd, Eng C. Biochemical screening and PTEN mutation analysis in individuals with autism spectrum disorders and macrocephaly. Eur J Hum Genet. 2014;22(2):273–276. doi:10.1038/ejhg.2013.114

66. Frazier TW. Autism Spectrum Disorder Associated with Germline Heterozygous PTEN Mutations. Cold Spring Harb Perspect Med. 2019;9(10). doi:10.1101/cshperspect.a037002

67. Götz M, Huttner WB. The cell biology of neurogenesis. Nat Rev Mol Cell Biol. 2005;6:777–788. doi:10.1038/nrm1739

68. Molyneaux BJ, Arlotta P, Menezes JRL, Macklis JD. Neuronal subtype specification in the cerebral cortex. Nat Rev Neurosci. 2007;8(June):427–437. doi:10.1038/nrn2151

69. Wegiel J, Flory M, Kuchna I, et al. Stereological study of the neuronal number and volume of 38 brain subdivisions of subjects diagnosed with autism reveals significant alterations restricted to the striatum, amygdala and cerebellum. Acta Neuropathol Commun. 2014;2:141. doi:10.1186/s40478-014-0141-7

70. Ecker C, Ronan L, Feng Y, et al. Intrinsic gray-matter connectivity of the brain in adults with autism spectrum disorder. Proc Natl Acad Sci. 2013;110(32):13222–13227. doi:10.1073/pnas.1221880110

71. Acevedo B, Aron E, Pospos S, Jessen D. The functional highly sensitive brain: a review of the brain circuits underlying sensory processing sensitivity and seemingly related disorders. Philos Trans R Soc Lond B Biol Sci. 2018;373(1744). doi:10.1098/rstb.2017.0161

72. Tomasi D, Volkow ND. Reduced Local and Increased Long-Range Functional Connectivity of the Thalamus in Autism Spectrum Disorder. Cereb Cortex. 2019;29(2):573–585. doi:10.1093/cercor/bhx340

73. Mighell TL, Thacker S, Fombonne E, Eng C, O’Roak BJ. An Integrated Deep-Mutational-Scanning Approach Provides Clinical Insights on PTEN Genotype-Phenotype Relationships. Am J Hum Genet. 2020;106(6):818–829. doi:10.1016/j.ajhg.2020.04.014

74. Barrows CM, McCabe MP, Chen H, Swann JW, Weston MC. PTEN Loss Increases the Connectivity of Fast Synaptic Motifs and Functional Connectivity in a Developing Hippocampal Network. J Neurosci. 2017;37(36):8595–8611. doi:10.1523/JNEUROSCI.0878-17.2017

75. Wickersham IR, Lyon DC, Barnard RJO, et al. Monosynaptic Restriction of Transsynaptic Tracing from Single, Genetically Targeted Neurons. Neuron. 2007;53:639–647. doi:10.1016/j.neuron.2007.01.033

76. Hafner G, Witte M, Guy J, et al. Mapping Brain-Wide Afferent Inputs of Parvalbumin-Expressing GABAergic Neurons in Barrel Cortex Reveals Local and Long-Range Circuit Motifs. Cell Rep. 2019;28(13):3450–3461.e8. doi:10.1016/j.celrep.2019.08.064

77. Nair A, Treiber JM, Shukla DK, Shih P, Muller R-A. Impaired thalamocortical connectivity in autism spectrum disorder: a study of functional and anatomical connectivity. Brain. 2013;136(Pt 6):1942–1955. doi:10.1093/brain/awt079

78. Woodward ND, Giraldo-Chica M, Rogers B, Cascio CJ. Thalamocortical dysconnectivity in autism spectrum disorder: An analysis of the Autism Brain Imaging Data Exchange. Biol psychiatry Cogn Neurosci neuroimaging. 2017;2(1):76–84. doi:10.1016/j.bpsc.2016.09.002

79. Zingg B, Dong H-W, Tao HW, Zhang LI. Input-output organization of the mouse claustrum. J Comp Neurol. 2018;526(15):2428–2443. doi:10.1002/cne.24502

80. Wimmer VC, Bruno RM, de Kock CPJ, Kuner T, Sakmann B. Dimensions of a projection column and architecture of VPM and POm axons in rat vibrissal cortex. Cereb Cortex. 2010;20(10):2265–2276. doi:10.1093/cercor/bhq068

81. Petersen CCH. The functional organization of the barrel cortex. Neuron. 2007;56(2):339–355. doi:10.1016/j.neuron.2007.09.017

82. Atasoy D, Aponte Y, Su HH, Sternson SM. A FLEX switch targets Channelrhodopsin-2 to multiple cell types for imaging and long-range circuit mapping. J Neurosci. 2008;28(28):7025–7030. doi:10.1523/JNEUROSCI.1954-08.2008

83. Ogawa S, Kwon C-H, Zhou J, Koovakkattu D, Parada LF, Sinton CM. A seizure-prone phenotype is associated with altered free-running rhythm in Pten mutant mice. Brain Res. 2007;1168:112–123. doi:10.1016/j.brainres.2007.06.074

84. Vorhees C V, Williams MT. Assessing spatial learning and memory in rodents. ILAR J. 2014;55(2):310–332. doi:10.1093/ilar/ilu013

85. Takahashi N, Oertner TG, Hegemann P, Larkum ME. Active cortical dendrites modulate perception. Science. 2016;354(6319):1587–1590. doi:10.1126/science.aah6066

86. Wong CW, Or PMY, Wang Y, et al. Identification of a PTEN mutation with reduced protein stability, phosphatase activity, and nuclear localization in Hong Kong patients with autistic features, neurodevelopmental delays, and macrocephaly. Autism Res. 2018;11(8):1098–1109. doi:10.1002/aur.1950

87. Ebner C, Ledderose J, Zolnik TA, et al. Optically Induced Calcium-Dependent Gene Activation and Labeling of Active Neurons Using CaMPARI and Cal-Light. Front Synaptic Neurosci. 2019;11:16. doi:10.3389/fnsyn.2019.00016

