## Supplementary material for "The impact of phosphorylated Pten at threonine 366 on cortical connectivity and behaviour"

**Supplementary Figures**

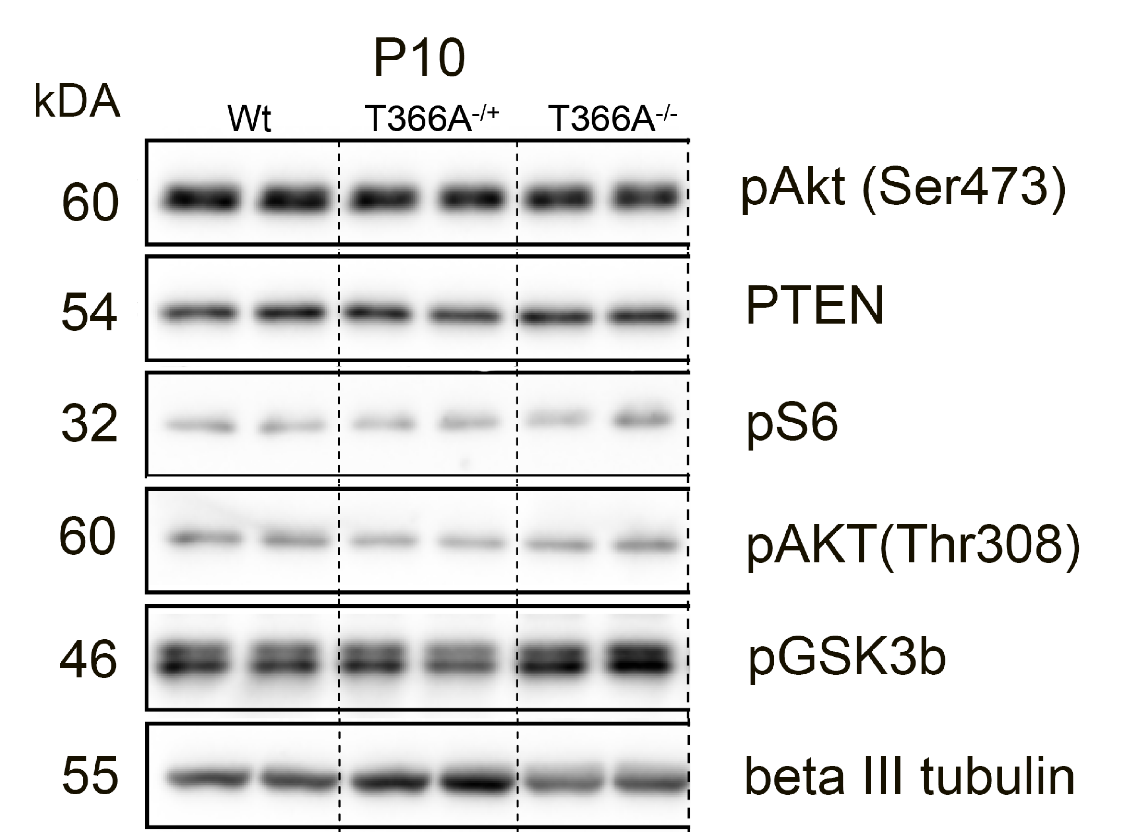

**Supplementary Figure 1: Protein levels in *Pten^T366A/T366A^* brains.**

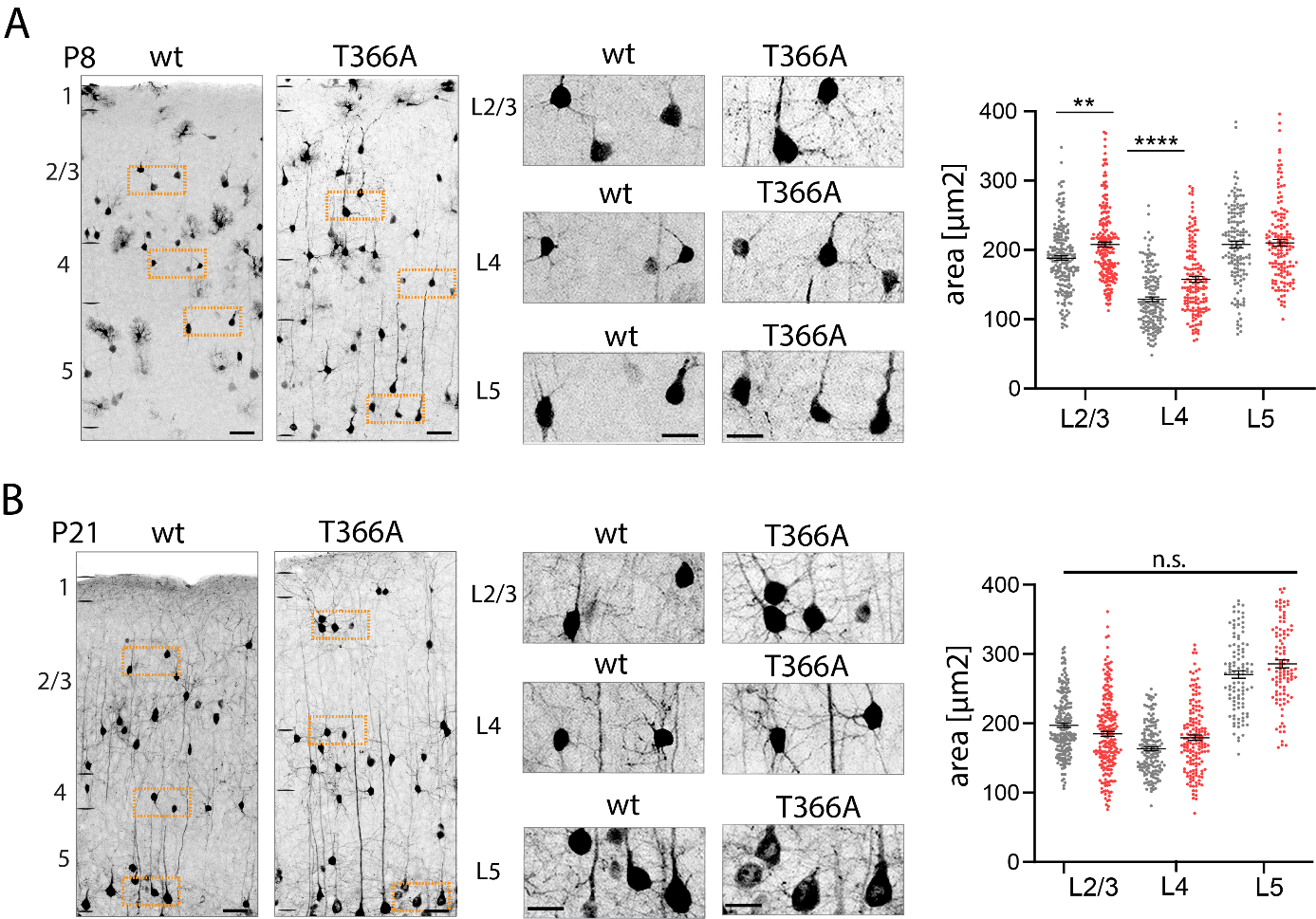

**Supplementary Figure 2: Soma size in *Pten^T366A/T366A^* cortical neurons at P8 and P21.**

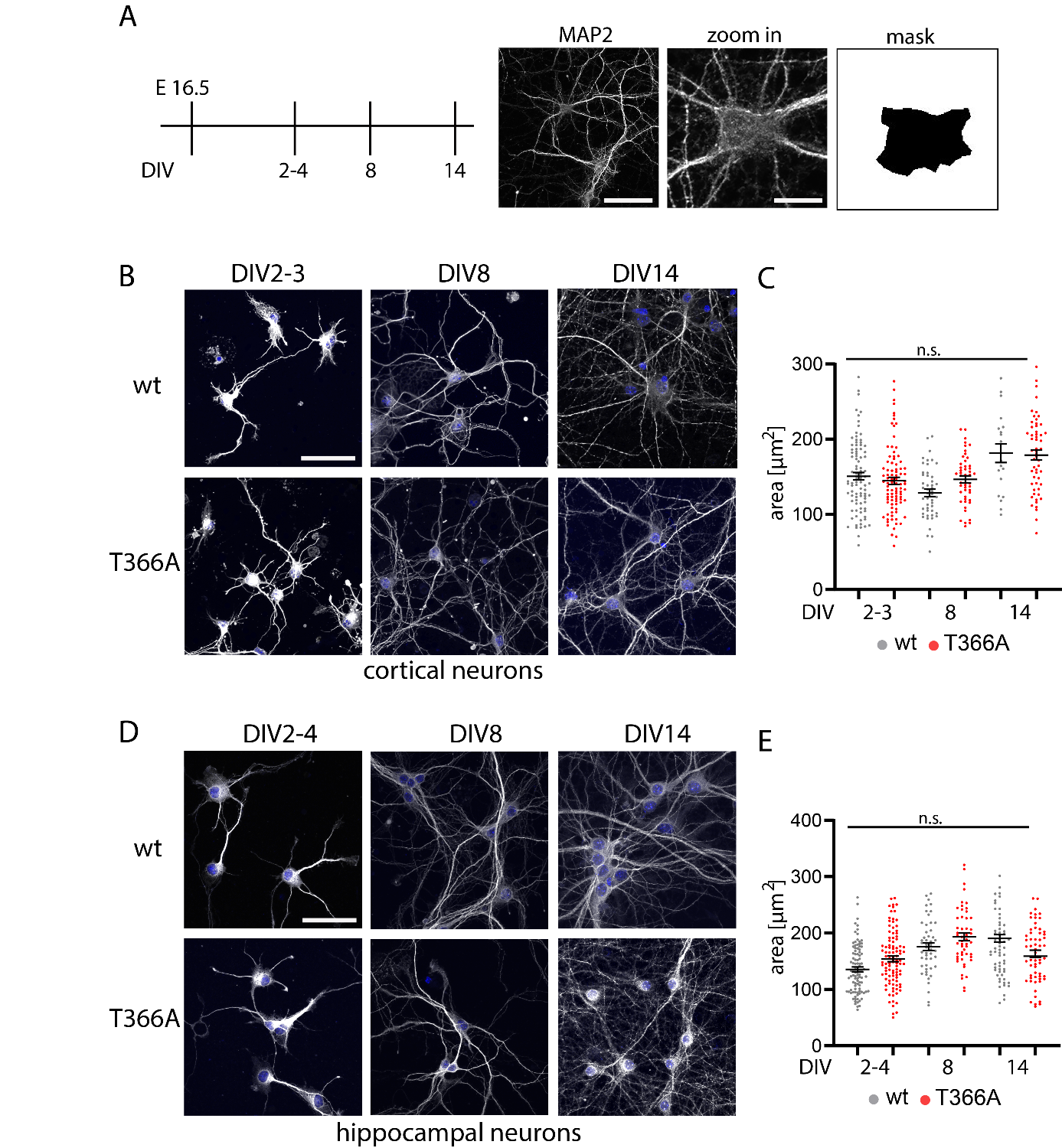

**Supplementary Figure 3: Soma size in *Pten^T366A/T366A^* primary cortical and hippocampal neurons.**

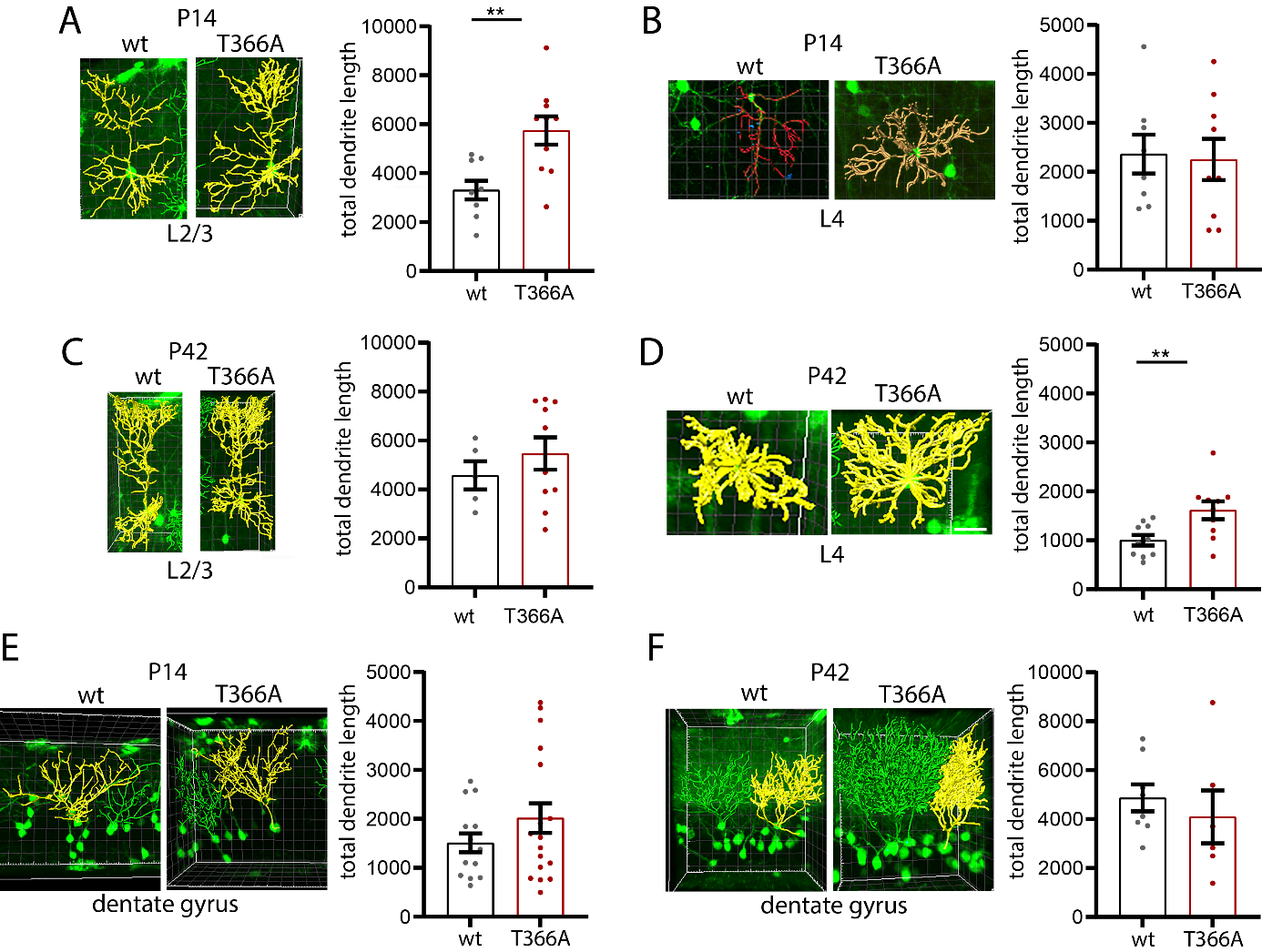

**Supplementary Figure 4: Dendritic lengths of cortical neurons in *Pten^T366A/T366A^* mice.**

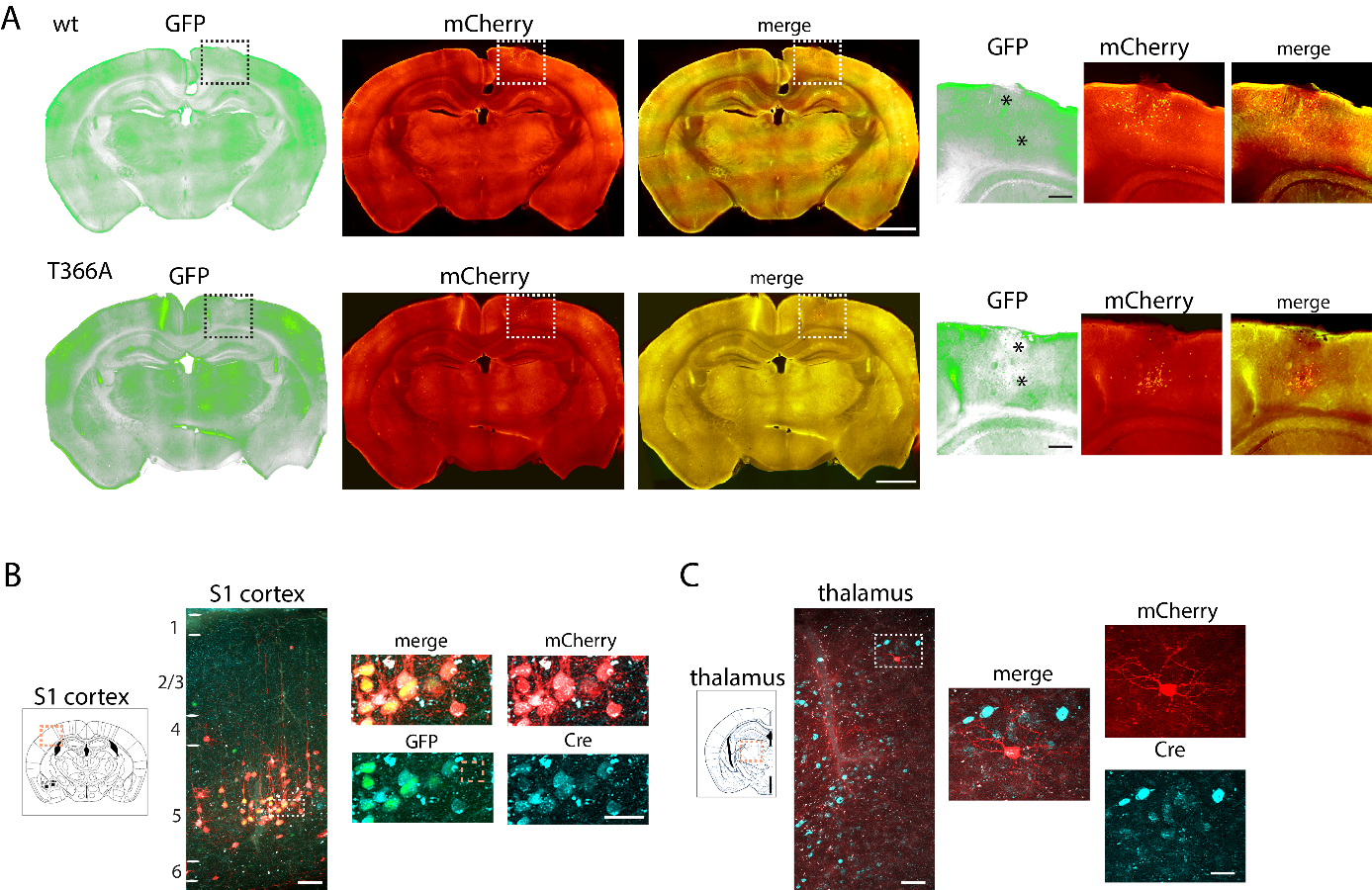

**Supplementary Figure 5: Injection sites and Cre expression in wt and in *Pten^T366A/T366A^* mice.**

**Supplementary Tables**

**Supplementary Table 1 Behaviour in *Pten^T366A/T366A^* and wt mice**

| **Genotype** | **Open field distance travelled (cm)** | **Open field center time (s)** | **Hanging wire** | **Vibrissae stimulation** | **grooming** | **rotarod** |  |
| --- | --- | --- | --- | --- | --- | --- | --- |
| **Wt** | 48.4±196.2 | 49.9±5.8 | 4.7±0.1 | 5.2±0.8 | 16.7 (mean) | 8.3±0.3 (day 1) | 8.9±0.6 (day 4) |
| ***Pten^T366A/T366A^*** | 48.2±207,1 | 52.0±6.4 | 4.7±0.1 | 1.5±0.5 | 12.3 (mean) | 14.0±0.9 (day 1) | 14.3±0.6 (day 4) |
| **P-value** | 0.9442 (n.s.) | 0.7342 (n.s.) | 0.9590 (n.s.) | 0.0002 *** | ^b^ >0.05 * | >0.9999 (n.s.) | >0.9999 (n.s.) |
|  | **Y maze** |  |  | **Barnes maze** |  | **Water maze** |  |
|  | *Successful alternations* | *Arm entries* | *Spontaneous alterations* | *Target quadrant* | *Other quadrants* | *Target quadrant* | *Other quadrants* |
| **Wt** | 18.1±1.3 | 29.4±2.1 | 62.7±2.2 | 40.2±25.4 | 19.9±1.8 | 44.4±4.7 | 18.5±1.6 |
| ***Pten^T366A/T366A^*** | 15.9±1.3 | 27.8±2.1 | 57.4±1.9 | 27.3±3.7 | 24.2±1.3 | 38.0±6.9 | 20.7±2.3 |
| **P-value^a^** | >0.9999 (n.s.) | >0.9999 (n.s.) | >0.9999 (n.s.) | 0.114 (n.s.), **^c^** 0.008 *** | >0.9999 (n.s.), **^c^** 0.0265 * | >0.9999 (n.s.), **^c^** 0.0006 *** | >0.9999 (n.s.), **^d^** 0.0024 ** |
|  | **Y maze male** |  |  | **Conditioned freezing males** |  |  |  |
|  | *Successful alternations* | *Arm entries* | *Spontaneous alterations* | *Habituation* | *Context* | *Pre-cues* | *cues* |
| **Wt** | 18.2±1.7 | 28.3±2.8 | 65.5±2.5 | 0.5±0.1 | 23.6±14.4 | 6.7±1.9 | 40.0±4.9 |
| ***Pten^T366A/T366A^*** | 13.0±1.5 | 25.1±2.8 | 52.4±1.7 | 0.7±0.1 | 16.8±4.0 | 8.8±2.4 | 41.2±5.7 |
| **P-value** | >0.9999 (n.s.) | >0.9999 (n.s.) | 0.0030 ** | >0.9999 (n.s.) | 0.0145 * | >0.9999 (n.s.) | >0.9999 (n.s.) |

Values are percentage mean (± standard error), data from 3 mice each genotype. Statistical analysis one-way ANOVA and unpaired t-test. P-value indicates significance level for comparison between, ^a^ wt and *Pten^T366A/T366A^* groups, ^b^ Analysis with Wilcoxon test, ^c^ between wt target – wt other quadrants, ^d^ between wt target *Pten^T366A/T366^* other quadrants.

**Supplementary Table 2 Cortical layer formation and proliferation of cortical progenitor neurons in *Pten^T366A/T366A^* and wt mice**

| **Genotype** | **L1 (%)** | **L2/3 (%)** | **L4 (%)** | **L5 (%)** | **L6 (%)** | **Neuron count** |
| --- | --- | --- | --- | --- | --- | --- |
| **Wt, NeuN** | 2.5±0.6 | 28.4±2.3 | 13.0±0.6 | 23.3±2.4 | 30.7±1.5 | 6293 NeuN |
| ***Pten^T366A/T366A^*, NeuN** | 2.5±0.5 | 32.7±1.5 | 14.5±2.3 | 21.9±1.3 | 28.2±2.4 | 8014 NeuN |
| **P-value** | >0.9999 (n.s.) | 0.4328 (n.s.) | >0.9999 (n.s.) | >0.9999 (n.s.) | >0.9999 (n.s.) |  |
| **Wt, Cux1** | - | 86.6±2.4 | - | - | - | 5282 Hoechst, 4486 Cux1 |
| ***Pten^T366A/T366A^*, Cux1** | - | 90.5±1.1 | - | - | - | 3517 Hoechst, 3172 Cux1 |
| **P-value** |  | 0.2107 (n.s) |  |  |  |  |
| **Wt, FoxP2** | - | - | - | - | 83.3±2.3 | 2324 Hoechst, 1928 FoxP2 |
| ***Pten^T366A/T366A^*, FoxP2** | - | - | - | - | 79.3±1.5 | 2769 Hoechst, 2196 FoxP2 |
| **P-value** |  |  |  |  | 0.1945 (n.s.) |  |
| **Wt, Ctip2** | - | - | - | 58.3±2.8 | - | 1822 Hoechst, 1070 CTIP2 |
| ***Pten^T366A/T366A^*, Ctip2** | - | - | - | 56.0±3.4 | - | 1345 Hoechst, 765 CTIP2 |
| **P-value** |  |  |  | 0.6056 (n.s.) |  |  |
| **E11/13-P1: wt** | 9.9±1.9 | 39.1±3.8 | 31.8±3.7 | 13.5±2.8 | 6.8±2.1 | 6837 BrdU |
| **E11/13-P1: *Pten^T366A/T366A^*** | 5.8±0.7 | 35.0±2.8 | 37.7±2.1 | 15.8±1.7 | 5.5±2.0 | 8864 BrdU |
| **P-value** | >0.9999 (n.s.) | >0.9999 (n.s.) | 0.3008 (n.s.) | >0.9999 (n.s.) | >0.9999 (n.s.) |  |
| **E13/15-P8: wt** | 2.1±0.5 | 64.8±4.1 | 20.4±3.6 | 9.2±1.3 | 3.3±0.9 | 7852 BrdU |
| **E13/15-P8: *Pten^T366A/T366A^*** | 1.5±0.7 | 58.1±4.2 | 19.4±3.8 | 7.2±1.0 | 9.4±3.3 | 5376 BrdU |
| **P-value** | >0.9999 (n.s.) | 0.4874 (n.s.) | >0.9999 (n.s.) | >0.9999 (n.s.) | 0.6381 (n.s.) |  |

Values are percentage mean (± standard error), data from 2 mice (layers), 3 mice (proliferation) each genotype. Statistical analysis two-way ANOVA for NeuN and proliferation, t-test for layer comparison. P-value indicates significance level for comparison between wt and *Pten^T366A/T366A^* groups.

**Supplementary Table 3 Soma size of cortical and hippocampal neurons in brain slices of *Pten^T366A/T366A^* and wt mice**

| **Genotype** | **L2/3 (µm^2^)** | **Neuron count** | **L4 (µm^2^)** | **Neuron count** | **L5 (µm^2^)** | **Neuron count** | **Dentate gyrus (µm^2^)** | **Neuron count** |
| --- | --- | --- | --- | --- | --- | --- | --- | --- |
| **P8, wt** | 188.2±3.3 | 203 | 128.8±3.4 | 155 | 207.9±4.7 | 146 | - | - |
| **P8, *Pten^T366A/T366A^*** | 207.8±3.7 | 203 | 157.5±4.1 | 155 | 210.1±4.7 | 146 | - | - |
| **P-value** | 0.0020 ** |  | <0.001 **** |  | >0.9999 (n.s.) |  |  | - |
| **P14, wt** | 163.5±1.9 |  | 131.9±2.3 |  | 238.4±4.7 | 181 | - | - |
| **P14, *Pten^T366A/T366A^*** | 195.4±2.1 |  | 173.9±2.9 |  | 147.7±4.7 | 181 | - | - |
| **P-value** | >0.0001 **** |  | >0.0001 **** |  | >0.9999 (n.s.) |  |  |  |
| **P21, wt** | 197.2±2.9 | 218 | 163.1±2.8 | 159 | 270.6±5.3 | 101 | - | - |
| **P21, *Pten^T366A/T366A^*** | 184.7±5.4 | 218 | 179.2±3.9 | 159 | 285.9±5.7 | 101 | - | - |
| **P-value** | 0.2102 (n.s.) |  | 0.0517 (n.s.) |  | 0.4076 (n.s.) |  |  |  |
| **P42 wt** | 131.1±1.7 | 319 | 91.4±2.2 | 146 | 253.4±4.7 | 197 | - | - |
| **P42, *Pten^T366A/T366A^*** | 130.0±1.9 | 319 | 95.6±2.2 | 146 | 241.7±4.3 | 225 | - | - |
| **P-value** | >0.9999 (n.s.) |  | >0.9999 (n.s.) |  | >0.1071 (n.s.) |  |  | - |
| **P14, wt** | - |  | - |  | - |  | 235.5±66.6 | 146 |
| **P14, *Pten^T366A/T366A^*** | - |  | - |  | - |  | 282.9±85.03 | 146 |
| **P-value** |  |  |  |  |  |  | >0.0001 **** |  |
| **P42, wt** | - |  | - |  | - |  | 106.2±29.2 | 253 |
| **P42, *Pten^T366A/T366A^*** | - |  | - |  | - |  | 97.3±26.5 | 366 |
| **P-value** |  |  |  |  |  |  | 0.0986 (n.s.) |  |

Values are percentage mean (± standard error), data from three mice each genotype. Statistical analysis with one-way ANOVA, p<0.05, unpaired t-test for data dentate gyrus, p<0.05. P-value indicates significance level for comparison between wt and *Pten^T366A/T366A^* groups.

**Supplementary Table 4 Soma size of cortical and hippocampal neurons in primary cell culture of *Pten^T366A/T366A^* and wt mice**

| **Genotype (neuron count)** | **2-4 DIV (µm^2^)** | **Neuron count** | **8 DIV (µm^2^)** | **Neuron count** | **14 DIV (µm^2^)** | **Neuron count** |
| --- | --- | --- | --- | --- | --- | --- |
| **Wt, cortical neurons** | 150.7±4.9 | 92 | 128.6±4.9 | 48 | 181.4±12.5 | 19 |
| ***Pten^T366A/T366A^*, cortical neurons** | 144.6±4.6 | 92 | 146.6±4.7 | 48 | 178.69±6.5 | 56 |
| **P-value** | >0.9999 (n.s.) |  | 0.6966 (n.s.) |  | >0.9999 (n.s.) |  |
| **Wt, hippocampal neurons** | 135.6±4.1 | 99 | 175.9±6.6 | 50 | 190.5±6.5 | 60 |
| ***Pten^T366A/T366A^*, hippocampal neurons** | 154.2±4.9 | 100 | 193.3±7.0 | 51 | 214.6±6.4 | 60 |
| **P-value** | 0.0923 (n.s.) |  | >0.9999 (n.s.) |  | 0.0904 (n.s.) |  |

Values are percentage mean (± standard error), data from two pregnant females each genotype, 5-8 embryos pooled. Statistical analysis with one-way ANOVA, p<0.05. P-value indicates significance level for comparison between wt and *Pten^T366A/T366A^* groups.

**Supplementary Table 5 Dendrite lengths of cortical and hippocampal neurons in *Pten^T366A/T366A^* and wt mice**

| **Genotype (neuron count)** | **L2/3 (µm^2^)** | **P-value** | **L4 (µm^2^)** | **P-value** | **Dentate gyrus (µm^2^)** | **P-value** |
| --- | --- | --- | --- | --- | --- | --- |
| **P14, wt (9)** | 3310±380.2 | 0.0032 ** | 2364±397.9 | 0.8523 (n.s.) | - |  |
| **P14, *Pten^T366A/T366A^* (10)** | 5742±578.2 | - | 2253±421.0 | - | - |  |
| **P42 wt (8)** | 4582±573.0 | 0.4024 (n.s.) | 1002±106.3 | 0.0097 ** | - |  |
| **P42, *Pten^T366A/T366A^* (10)** | 5471±659.0 | - | 1614±182.8 |  | - | - |
| **P14, wt (10)** | - | - | - |  | 1509±191.9 | 0.1947 (n.s.) |
| **P14, *Pten^T366A/T366A^* (18)** | - | - | - |  | 2018±302.6 | - |
| **P42 wt (8)** | - | - | - |  | 4870±550.7 | 0.5031 (n.s.) |
| **P42, *Pten^T366A/T36A6^* (6)** | - | - | - |  | 4092±108.3 | - |

Values are percentage mean (± standard error), data from three mice each genotype. Statistical analysis with unpaired t-test, p< 0.001. P-value indicates significance level for comparison between wt and *Pten^T366A/T366A^* groups of one age each.

**Supplementary Table 6 Presynaptic input to S1 cortex in *Pten^T366A/T366A^* and wt mice**

| **Genotype** | **Local S1** | **Long-range** | **Visual cortices** | **S2 cortex** | **Motor cortices** | **thalamus** | **contralateral S1** |
| --- | --- | --- | --- | --- | --- | --- | --- |
| **Wt %** | 73.6±3.4 | 26.3±3.4 | 18.0±1.6 | 4.6±2.3 | 38.9±4.6 | 21.4±2.0 | 16.9±9.8 |
| **Wt, neuron count** | 1809 | 611 | 126 | 22 | 250 | 134 | 79 |
| ***Pten^T366A/T366A^* %** | 71.6±10.2 | 28.3±10.2 | 12.0±6.4 | 3.3±2.5 | 15.9±4.6 | 46.9±3.8 | 21.8±7.3 |
| ***Pten^T366A/T366A^* , neuron count** | 1071 | 327 | 25 | 10 | 42 | 162 | 88 |
| **P-value** | 0.8591 (n.s.) | 0.8591 (n.s.) | 0.5118 (n.s.) | 0.7288 (n.s.) | 0.0245 * | 0.0042 ** | 0.7108 (n.s.) |

Values are percentage mean (± standard error), data from three mice each genotype. Statistical analysis with unpaired t-test, p< 0.05. P-value indicates significance level for comparison between wt and *Pten^T366A/T366A^* groups
